# Translational activators align mRNAs at the small mitoribosomal subunit for translation initiation

**DOI:** 10.1101/2025.01.26.634913

**Authors:** Joseph B. Bridgers, Andreas Carlström, Dawafuti Sherpa, Mary T. Couvillion, Urška Rovšnik, Jingjing Gao, Bowen Wan, Sichen Shao, Martin Ott, L. Stirling Churchman

## Abstract

Mitochondrial gene expression is essential for oxidative phosphorylation. Mitochondrial-encoded mRNAs are translated by dedicated mitochondrial ribosomes (mitoribosomes), whose regulation remains elusive. In the baker’s yeast *Saccharomyces cerevisiae*, nuclear-encoded mitochondrial translational activators (TAs) facilitate transcript-specific translation by a yet unknown mechanism. Here, we investigated the function of TAs containing RNA-binding pentatricopeptide repeats (PPRs) using selective mitoribosome profiling and cryo-EM structural analysis. These analyses revealed that TAs exhibit strong selectivity for mitoribosomes initiating on their target transcripts. Moreover, TA-mitoribosome footprints indicated that TAs recruit mitoribosomes proximal to the start codon. Two cryo-EM structures of mRNA-TA complexes bound to post-initiation/pre-elongation-stalled mitoribosomes revealed the general mechanism of TA action. Specifically, the TAs bind to structural elements in the 5’ UTR of the client mRNA as well as to the mRNA channel exit to align the mRNA in the small subunit during initiation. Our findings provide a mechanistic basis for understanding how mitochondria achieve transcript-specific translation initiation without relying on general sequence elements to position mitoribosomes at start codons.

## Introduction

Mitochondria derive from an alpha-proteobacteria ancestor and have retained a reduced genome, which encodes critical components of the oxidative phosphorylation (OXPHOS) complexes. The mitochondrial genome of *Saccharomyces cerevisiae*, or budding yeast, encodes seven OXPHOS proteins and one component of the mitochondrial ribosome (mitoribosome). These genes are transcribed by a dedicated mitochondrial RNA polymerase and translated by specialized mitoribosomes^1^. The remaining thirty-eight OXPHOS subunits are transcribed from the nuclear genome and are synthesized on cytoplasmic ribosomes prior to their import into mitochondria. Translation of nuclear and mitochondrial subunits belonging to the same OXPHOS complex is coordinated to prevent the production of excess, unassembled subunits that can be detrimental to the cell through the generation of reactive oxygen species and chronic proteostatic stress^2–6^.

Despite sharing some similarities with their bacterial ancestors, mitoribosomes have diverged significantly from bacterial ribosomes, as evidenced by substantially differing structures and mechanisms of translation. Bacterial translation initiation often employs the Shine-Dalgarno (SD) sequence motif, which is found in the 5’ untranslated region (5’ UTR) of mRNAs just upstream of the start codon. The SD motif base pairs with the anti-SD sequence in the 16*S* rRNA, which helps to recruit the mRNA to the small subunit (SSU). The distance between the SD sequence and the start codon is important to align the start codon with the ribosomal P-site to ensure fidelity and efficiency of translation initiation^7^. How translation of mitochondrial mRNAs is initiated is yet unknown because mRNAs in mitochondria lack SD-like motifs, and the mitoribosome lacks an anti-SD sequence.

Early genetic work in yeast identified a group of nuclear-encoded mitochondrial proteins that are necessary for translation of individual mitochondrial transcripts and were termed translational activators (TAs)^6,8^. TAs have been proposed to recruit their target mRNAs to the mitoribosome and aid in translation initiation^6,8,9^. A subset of the TAs also participates in translation feedback loops to control synthesis of specific proteins in relation to the efficiency by which they can assemble into OXPHOS complexes^10–13^. TA binding to mRNAs is supported by specific helical structural motifs, including pentatricopeptide repeats (PPRs), which have been proposed to mediate sequence-specific mRNA binding^14–20^. Several studies revealed functional genetic interactions between the TAs and the 5’ UTRs of their target transcripts, but only a few of the PPR-containing TAs have been shown to directly bind RNA^21–24^. Moreover, genetic and protein proximity mapping revealed that TAs bind to the mRNA channel exit (MCE) of the SSU^9,25^, which would be in line with a role in translation initiation. While it is clear that TAs are required for the translation of specific transcripts within mitochondria, how TAs engage with the 5’ UTR or the mitoribosome to support translation initiation and elongation remains unresolved.

In this study, we determined when and how TAs engage with the mitoribosome during translation. By establishing selective ribosome profiling for mitoribosomes (sel-mitoRP), we determined when TAs bind the mitoribosome on target transcripts. These analyses revealed that TAs are highly enriched on mitoribosomes near the translation initiation site of their client mRNA and that they are released soon after the onset of elongation, confirming a role in initiation. Moreover, TA binding patterns in the 5’ UTR suggest the stabilization of specific mRNA folds by complex formation between this portion of the mRNA and the TAs. Consequently, inhibiting translation elongation by genetic ablation of a general elongation factor led to the accumulation of post-initiation/pre-elongation mitoribosomes with bound TAs. Structure determination using single-particle cryogenic electron microscopy (cryo-EM) resolved two distinct TA-mitoribosome complexes with bound mRNA. The structures revealed how the *ATP8* and the *ATP9* TAs bind to the mitoribosome, with the 5’ UTR of their client mRNA wrapped around the TA(s) and the start codon base-paired with the initiator tRNA at the P-site. Additional structural analyses of *ATP9*-TA complexes affinity purified with elongation-competent mitoribosomes further corroborated the role of TAs specifically in translation initiation. Together our data demonstrate that TAs function to recruit their target transcript to the mitoribosome and to act as molecular guide to position the transcript in the mRNA channel for proper translation initiation.

## Results

### Interaction of translational activators with the mitochondrial translatome

The RNA binding capabilities of the PPR proteins as well as their localization to the mRNA channel exit (MCE) position them as candidate factors for serving an SD-like function in recruiting their target transcript to the mitoribosome. To investigate the molecular functions of TAs containing PPR motifs during mitochondrial translation, we adapted selective ribosome profiling for mitochondrial ribosomes (sel-mitoRP) (Fig. 1a,b and Extended Data Fig. 1, 2)^26,27^. This method determines when specific factors interact with the mitoribosome at codon resolution. Specifically, lysate from crosslinked cells was treated with RNAse and total ribosomes were isolated. Mitoribosome subpopulations bound by a TA of interest were purified by affinity chromatography and the ribosome-protected RNA fragments were analyzed by sequencing (Fig. 1b). We first performed sel-mitoRP for Pet111, the TA of *COX2*^28^, and for Mrps17, a constitutive subunit of the mitoribosome. We have previously shown that mitoribosome footprint density on the *COX2* transcript is lost upon deletion of *PET111*^2^. Consequently, sel-mitoRP analyses demonstrated a strong and specific enrichment of Pet111-mitoribosome footprints on *COX2*, with the highest enrichment of RNA footprints in the 5’ UTR, approximately 200 times that of the total mitoribosome control (Fig. 1c,d and Extended Data Fig. 3). The enrichment of Pet111-mitoribosome footprints just upstream of the start codon aligns with previous studies which have shown the importance of the *COX2* 5’ UTR in Pet111 translational activation^23,29^.

**Figure 1:**
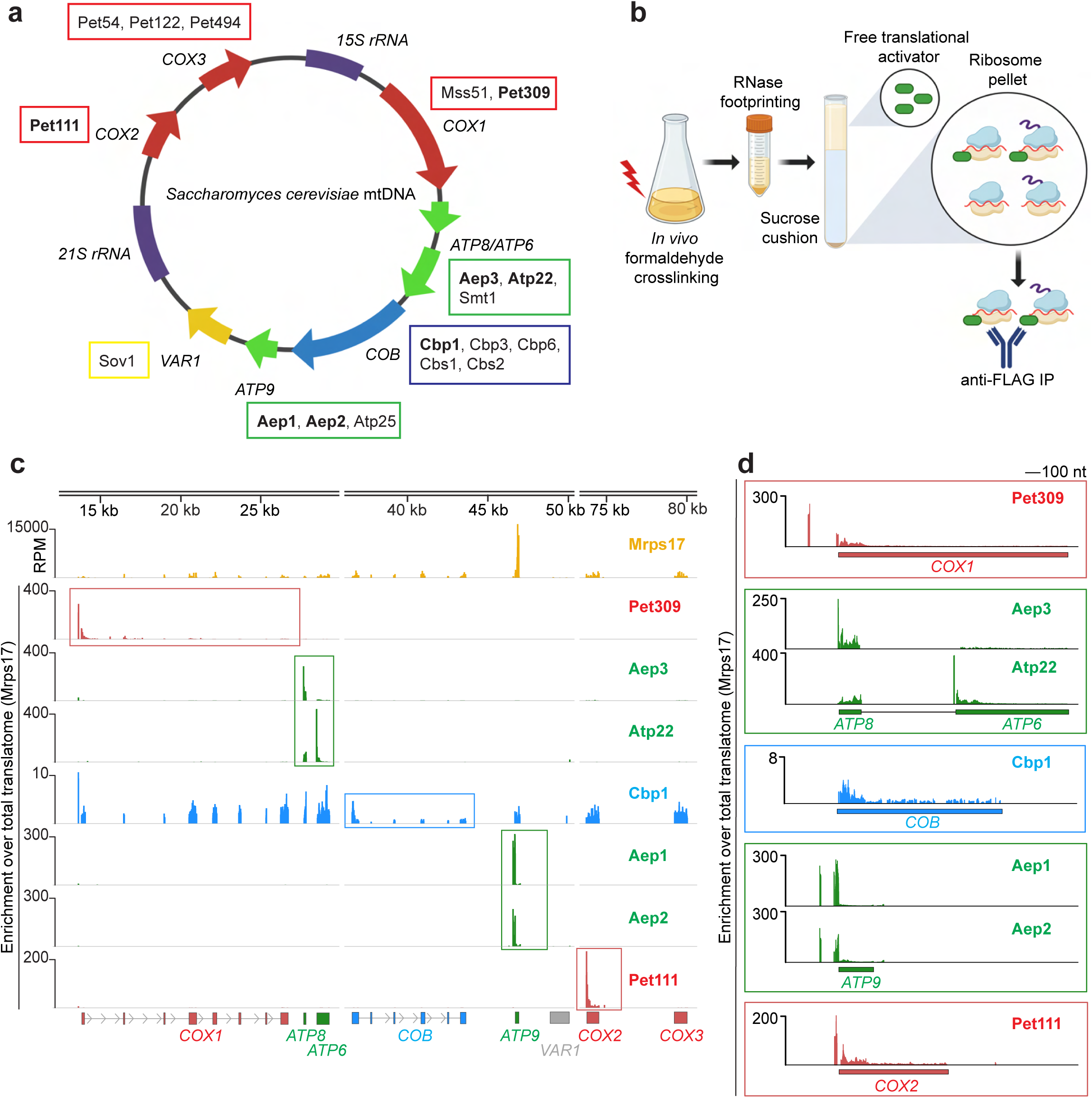
PPR domain TAs are selectively enriched at translation initiation. **a,** Schematic of *S. cerevisiae* mitochondrial genome. The locations of the 15S and 21S rRNAs are labeled in purple and the protein coding genes are color-coded based on the complex they belong to: ATP synthase in green, Complex III in blue, and Complex IV in red. Translational regulators are boxed in the corresponding color next to their target transcript. Proteins that are analyzed in panel c are highlighted in bold. **b,** Overview of selective mitoribosome profiling (sel-mitoRP). **c,** Top of panel: Inferred A-site counts for a total mitoribosome control dataset (Mrps17). RPM = reads per million mapped reads. Bottom of panel: Sel-mitoRP of the indicated TA normalized to the total mitoribosome control (Mrps17). The boxed genes are the known translation target of the given TA. **d,** Zoom in on the enrichment for the given TA on the boxed target gene from c. All data consist of combined reads from at least two biological replicates.

To determine if the other mitochondrial TAs are as specific as Pet111, we performed sel-mitoRP for TAs containing PPR motifs: Pet309 (TA for *COX1*), Aep3 (TA for *ATP8*), Atp22 (TA for *ATP6*), Cbp1 (TA for *COB*), and Aep1 and Aep2 (TAs for *ATP9*) (Fig. 1c,d and Extended Data Fig. 3)^15–20^. Similar to Pet111, the majority of the TAs showed a strong enrichment on mitoribosomes translating their target transcript (Fig. 1c). The only exception to this trend was Cbp1, which did not show a specific enrichment on the *COB* transcript, which may be explained by its role as part of a translational feedback loop^13^.

### TA-mitoribosome footprints are enriched upstream of start codons

Closer examination of the target transcript for each TA indicated a strong enrichment of TA-mitoribosome footprints in the 5’ UTR, decreasing abruptly after the start codon and tailing off further after early elongation (Fig. 1d). This enrichment at the 5’ end of open reading frames and depletion from the 3’ end argues that the TAs leave the ribosome soon after initiation. To gain more insights into how TAs influence initiation, we performed a closer analysis of the footprints around translation start sites as the locations and heterogeneity of footprint ends shed light on the state of the ribosome^30–32^.

We began with footprints of *ATP9* mRNA, which are highly abundant (Fig. 1c) due to the high rates of Atp9 synthesis required to support assembly of the ATP synthase rotor that contains ten Atp9 subunits. Comparing footprints from Mrps17-sel-mitoRP with and without formaldehyde crosslinking revealed different footprints for the initiating mitoribosome. The initiating mitoribosome on *ATP9* in non-crosslinked mitoribosome profiling yielded a footprint with 5’ ends 16 nucleotides upstream of the start codon and 3’ ends approximately 22 nucleotides downstream of the start codon (Fig. 2a), in line with the expected size and position of a mitoribosome-protected fragment with the start codon positioned in the P site. Interestingly, crosslinking stabilized a larger footprint in the 5’ UTR with 5’ ends approximately 38 nucleotides upstream of the start codon and variable 3’ ends within a few nucleotides of the start codon (Fig. 2a). This upstream footprint was also highly enriched in sel-mitoRP for the *ATP9* TAs, Aep1 and Aep2 (Fig. 2a). Due to an empty A-site of the initiating ribosome, cytosolic ribosomes can have a truncated 3’ footprint end near the start codon via RNase I cleavage inside the ribosome^30–32^. Therefore, we hypothesized that the larger upstream footprint might reflect an initiating TA-mitoribosome complex with the TAs, extending the 5’ end of the mitoribosome footprint through increased protection of the mRNA. Additionally, the 3’ end of the footprint is truncated from the expected +22 position, likely due to RNase I cutting within the initiating mitoribosome. Without crosslinking to stabilize the TAs, footprints due to RNase I internal cleavage would be too small to be sequenced, which is likely why they were not observed in the natively prepared library.

**Figure 2:**
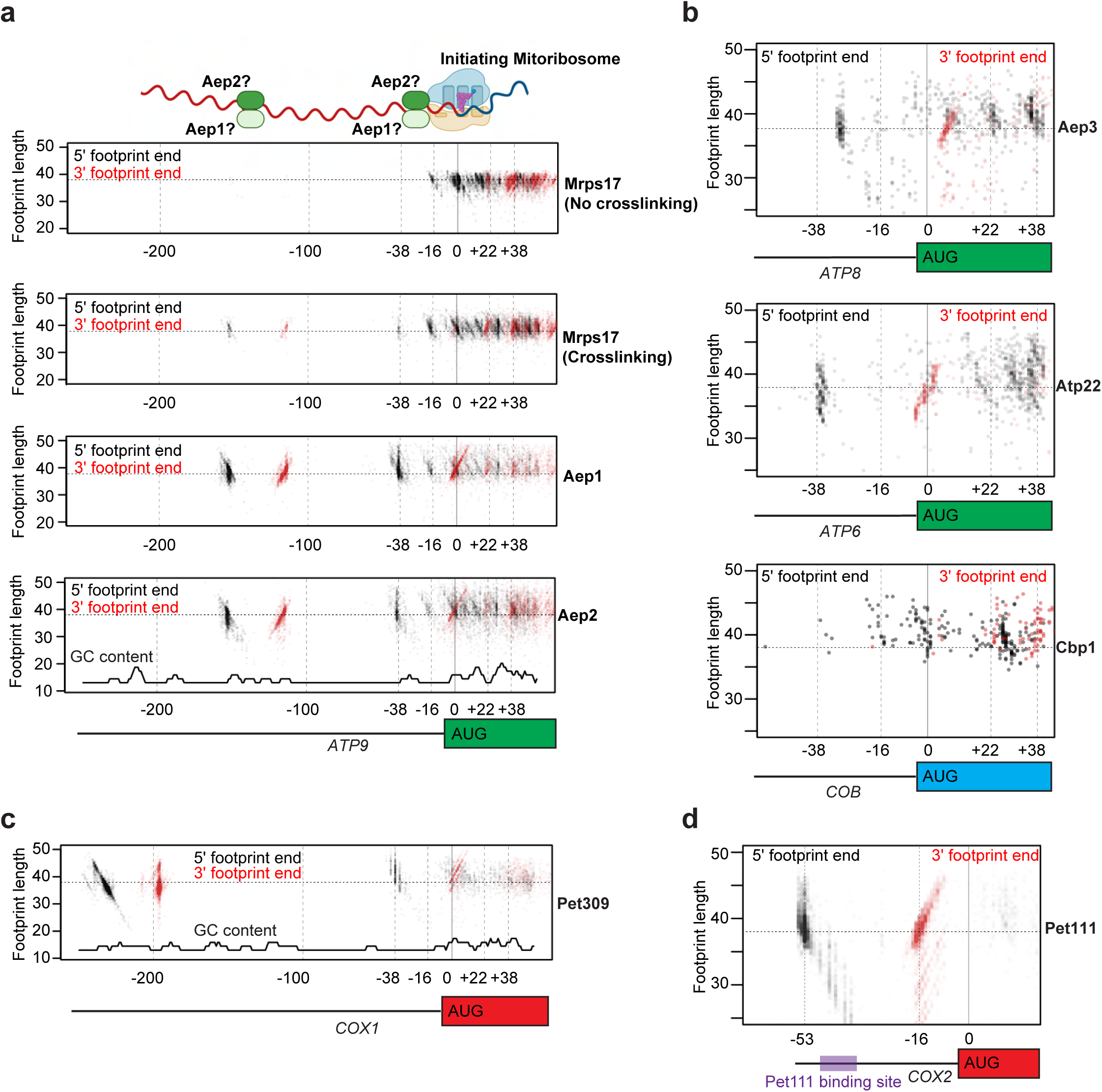
Visualizing TA-mitoribosome footprints in the 5’ UTR. **a,** The 5’ and 3’ ends of footprints from total mitoribosome profiling (Mrps17) either with or without formaldehyde crosslinking were plotted in relation to the start codon of *ATP9* and compared to the footprints from Aep1 and Aep2 sel-mitoRP. The horizontal dashed line indicates a fragment length of 38 nucleotides, which is the approximate size of a mitoribosome footprint at initiation. GC content of the *ATP9* transcript is plotted along the bottom of the Aep2 footprint data. The diagram at the top of the panel depicts the position of an initiating mitoribosome footprint with its 5’ end at -16 and its 3’ end at +22. Aep1 and Aep2 are depicted upstream of the initiating mitoribosome to indicate the potential binding sites of Aep1 or Aep2 in the 5’ UTR and the subsequent extension of the 5’ end from -16 to -38. **b,** 5’ and 3’ footprint ends for each sel-mitoRP dataset with footprints that abut the start codon, indicating a potential TA-mitoribosome complex at initiation. **c,** 5’ and 3’ footprint ends from Pet309 sel-mitoRP plotted around the start codon of *COX1*. GC content of the transcript is plotted as a solid black line at the bottom of the panel. **d,** 5’ and 3’ footprint ends from Pet111 sel-mitoRP plotted around the start codon of *COX2*. The known Pet111 binding site in the 5’ UTR of *COX2* is boxed in purple. All data consist of combined reads from at least two biological replicates.

We next analyzed the 5’ and 3’ ends of the footprints from sel-mitoRP for the other PPR proteins to determine if they had patterns similar to Aep1 and Aep2 in the 5’ UTR of their target transcripts (Fig. 2b,c,d and Extended Data Fig. 4). Sel-mitoRP data for Atp22 and Pet309, the TAs for *ATP6* and *COX1* respectively, exhibited footprint ends that were very similar to those for Aep1 and Aep2 (Fig. 2b,c). Aep3-mitoribosomes yielded a 3’-shifted RNA footprint, presumably because Aep3 protects a shorter ribosome-adjacent stretch of RNA, and the Cbp1 footprint in the *COB* UTR was less intense compared to the downstream initiating footprint, perhaps due to its reduced enrichment on its target transcript.

The Pet111 sel-mitoRP footprints had a unique pattern in the 5’ UTR with well-defined 5’ ends positioned 53 nucleotides upstream of the start codon and more variable 3’ ends positioned around 16 nucleotides upstream of the start codon (Fig. 2d and Extended Data Fig. 4). This correlates well with the experimentally defined Pet111 binding site in the 5’ UTR of *COX2*^23^ and an AlphaFold3 model of Pet111 bound to the *COX2* 5’ UTR (Extended Data Fig. 5). All these TA-mitoribosome footprints suggest that they could function similarly to Aep1 and Aep2 by binding to the mitoribosome and upstream RNA to position the mitoribosome at the start codon.

In addition to the TA-mitoribosome footprints proximal to the start codon, Aep1, Aep2, and Pet309 sel-mitoRP revealed RNase protected fragments further upstream in the 5’ UTR (Fig 2a,c). Specifically, a second set of mitoribosome footprints for Aep1 and Aep2 sel-mitoRP was found over 100 nucleotides upstream of the *ATP9* start codon, and Pet309 sel-mitoRP upstream footprints were more than 200 nucleotides upstream of the *COX1* start codon (Fig. 2a,c). These footprints could represent a separate population of mitoribosomes or they could be RNA fragments interacting with the initiating mitoribosome through looping and folding of the 5’ UTR. Although the patterning and positioning of the RNA footprints differed across the TAs studied, in all cases the TAs engaged with the mitoribosome proximal to the start codon of their target transcript and were disenriched during early elongation, suggesting a role in RNA recruitment to the mitoribosome during translation initiation.

### Cryo-EM structures of initiating mitoribosomes

Sel-mitoRP revealed that TAs bind to mitoribosomes transiently during early stages of translation through the formation of specific contacts with the mitoribosomes and their client mRNA. Inspired by these results and to gain a more detailed perspective, we aimed to stall mitoribosomes at initiation to determine their structures by single particle cryogenic electron microscopy (cryo-EM). We genetically induced initiation-stalled mitoribosomes by deleting *TUF1*, coding for the homolog of the bacterial translation elongation factor EF-Tu. Without Tuf1, elongator tRNAs cannot be delivered to the A-site, hence mitoribosomes are trapped at a post-initiation/pre-elongation stage with an initiator fMet-tRNA in the P-site^33^ (Fig. 3a). We enriched for these stalled mitoribosome complexes through centrifugation and performed single-particle cryo-EM (Fig. 3b). After initial data processing (Extended Data Fig. 6), focused classification on the subunit interface of mitochondrial monosomes revealed absence of tRNA in the A-site, with a single tRNA found in either the P-or E-site, which is in line with mitoribosomes unable to elongate (Fig. 3c and Extended Data Fig. 7a). After further focused refinement and classification on the SSU, we observed two major classes of mitoribosomes with additional mRNA and protein densities containing clear signatures of PPR domains located at the MCE (Fig. 3d,e and Extended Data Figs. 6c and 7b,d). Rigid-body fitting of AlphaFold2-predicted structures of PPR-containing TAs into the extra densities identified the factors bound to these two post-initiation/pre-elongation complexes as Aep3 (Fig. 3d), the TA for *ATP8*, and a complex of Aep1, Aep2 and Atp25C (Fig. 3e), the TAs for *ATP9*.

**Figure 3:**
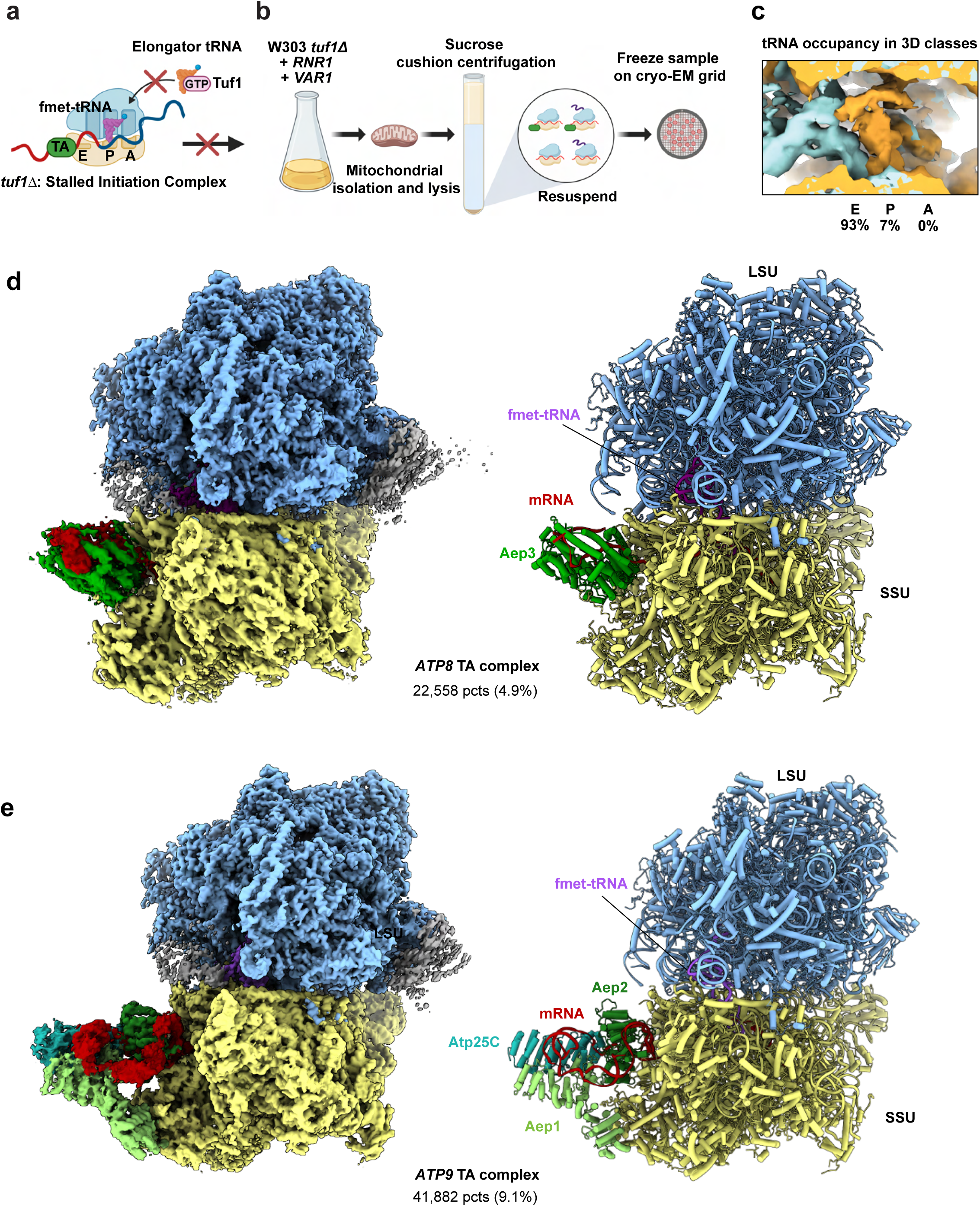
*ATP9* and *ATP8* TAs bind mitoribosomes stalled at a post-initiation, pre-elongation stage. **a,** Schematic of *TUF1* deletion which leads to stalling of mitoribosomes at initiation by preventing the recruitment of elongator tRNAs. **b,** Overview of sample preparation of *tuf1Δ* strain mitoribosomes for cryo-EM analysis. **c,** Occupancy of tRNA densities in the decoding center on the SSU obtained after focused 3D classification on 461,825 monosome particles. 93% of classes showed a tRNA density in the E-site and 7% in the P-site. No density could be visualized in the A-site in any of the classes, which is indicative of ribosomes able to initiate translation but not proceed to elongation**. d,** Cryo-EM density map (left) and structure model (right) of *ATP8* TA, Aep3 (green), bound to the small subunit of the mitoribosome. **e,** Cryo-EM density map (left) and structure model (right) of the *ATP9* TAs Aep1 (light green), Aep2 (green), and Atp25C (cyan) bound to the small subunit of the mitoribosome.

### Aep3 engages the initiating mitoribosome

Aep3 is predicted to consist of an array of PPR motifs that overall could be rigid-body fitted into the extra helical densities that generally sit perpendicular to the mitoribosome in the cryo-EM map at an overall resolution of 3.7 Å (Figs. 3d and 4). However, some loops were less well resolved, and unstructured N-terminal (residues 33 to 52) and C-terminal (residues 531 to 558) segments were not observed in the density map and hence were not modeled. Aep3 makes direct contacts with the MCE through mitoribosomal protein Mrp51 (bS1m), which has been genetically linked to the *COX2* and *COX3* 5’ UTRs^34^ (Fig. 4c). Interestingly, an extended density for bS1m, which was not resolved in other mitoribosome structures, was observed here and presumably forms a contact and binding platform for Aep3 (Fig. 4c,d). A density representing the *ATP8* mRNA was detected, extending from the P-site bound tRNA to the MCE (Fig. 4a,b). The 5’ UTR of *ATP8* mRNA interacts with a positively charged RNA binding pocket of Aep3 that is formed by a helical array of its PPR motifs (Fig. 4d-f). An RNA stem loop with three branches is predicted to form just upstream of where the 5’ end of the Aep3-mitoribosome footprint lies (Fig. 2b and Extended Data Fig. 8a). Some of this predicted secondary structure might explain the mRNA density occupying the Aep3 RNA pocket (Fig 4e); however the broken mRNA density cannot resolve the precise sequence of the RNA. The extensive RNA contacts made by the Aep3 PPR motifs at the MCE is in line with the additional RNAse-protection of the *ATP8* 5’ UTR upstream of the initiating mitoribosome in the Aep3 sel-mitoRP data (Fig. 2b and Extended Data Fig. 4b). These data confirm and expand on the results from Aep3 sel-mitoRP that Aep3 binds to the 5’ UTR of its target transcript and to the MCE, presumably to position *ATP8* mRNA for translation initiation.

**Figure 4:**
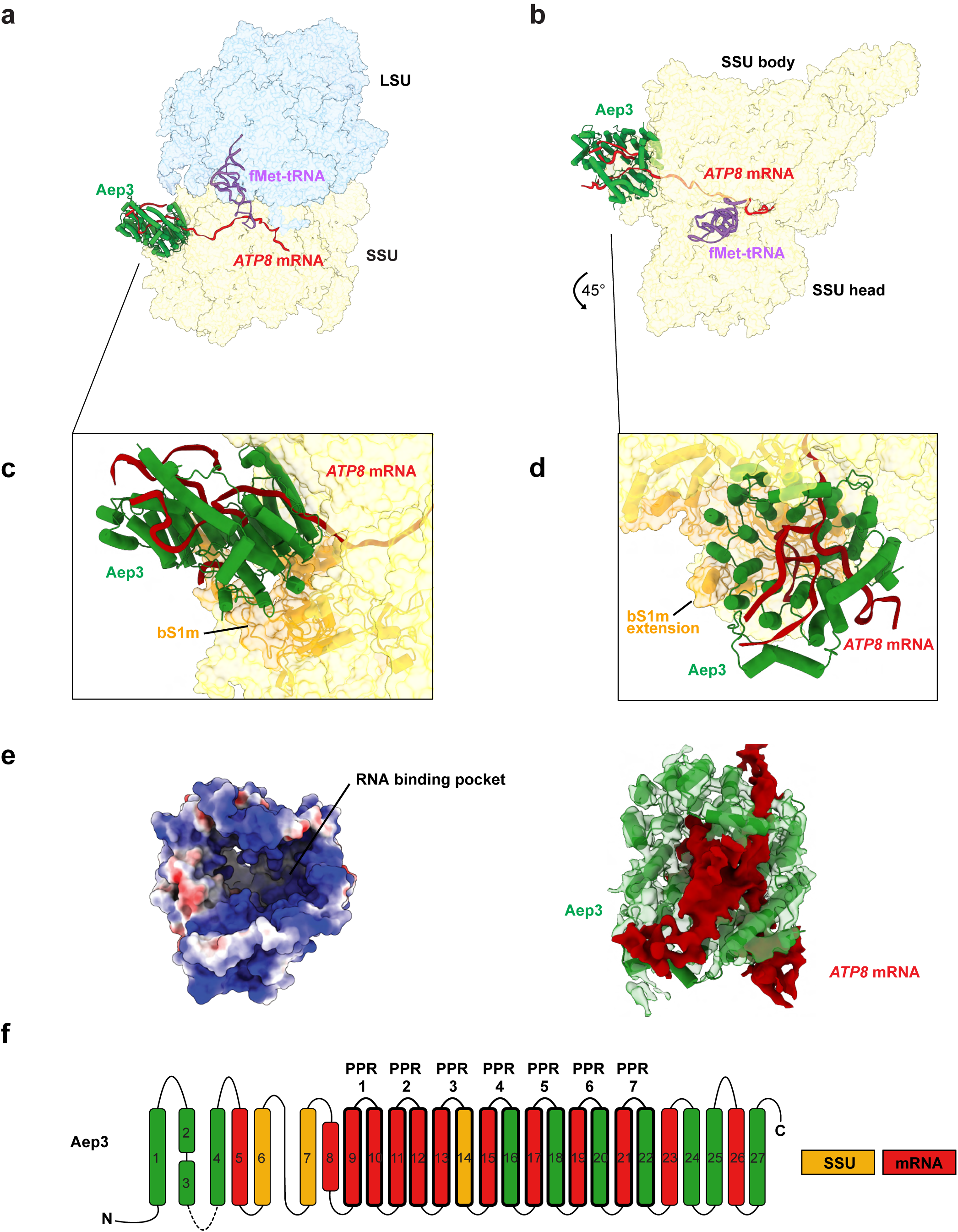
Structure of Aep3 bound to the initiating mitoribosome and 5’ UTR of *ATP8*. **a,** Side view of Aep3 (green) engaging with the mitoribosome SSU (yellow) and the *ATP8* mRNA. **b,** Top view demonstrating the interaction of the *ATP8* mRNA with Aep3 PPR motifs. **c,** Zoom in from panel a highlighting interaction of Aep3 with the SSU protein Mrp51 (bS1m) and the extended density that forms a binding platform. **d,** Zoom in from panel b demonstrating the RNA density that is surrounded by Aep3 PPR motifs. **e,** (Left) Aep3 surface map of PPR motifs colored by charge with positively charged residues in blue and negatively charged residues in red. (Right) Aep3 PPR motifs overlayed with the *ATP8* mRNA density. **f,** Linear diagram of Aep3 demonstrating which PPR motifs and alpha-helices are engaged with the SSU (yellow) or the *ATP8* mRNA (red). The dashed line represents residues 33 to 52 which were not resolved in the structure.

### *ATP9* TAs bind mitoribosomes at initiation

The other class, obtained from 9.1% of the particles from the dataset, contained a much larger complex bound to the MCE (Fig. 3e and Extended Data Figs. 6c and 7b,c). Using AlphaFold2 and AlphaFold3, we predicted and rigid-body fitted the individual structures of the three *ATP9* TAs, Aep1, Aep2 and Atp25 into the TA density of this map at a resolution of 3.8 Å (Fig 5a,b and Extended Data Fig. 9). The existence of a single complex containing both Aep1 and Aep2 explains the highly similar sel-mitoRP profiles of Aep1 and Aep2 (Figs. 1d and 2a), as well as their co-precipitation from lysate (Fig. 5e). Each of these proteins demonstrated a predicted overall conformation that fitted well into defined parts of the density^35^ (Figs. 3e, 5a,b, and Extended Data Fig. 9). For Aep1, the majority of the protein could be fit with the exception of the N-terminal residues 1-41, corresponding to the predicted mitochondrial targeting signal (MTS). For Aep2, residues 1-94 of the N-terminus and residues 536-580 of the C-terminus did not have a corresponding map density and were therefore not modeled. Interestingly, Atp25 was previously shown to be processed into an N-terminal protein of 29 kDa and a C-terminal protein of 36 kDa that has a specific role in *ATP9* translation^36^. This C-terminal part of Atp25 (Atp25C), ranging between residues 296 and 609, fit well into our cryo-EM map, which is consistent with its proposed role in *ATP9* mRNA stabilization^37^.

**Figure 5:**
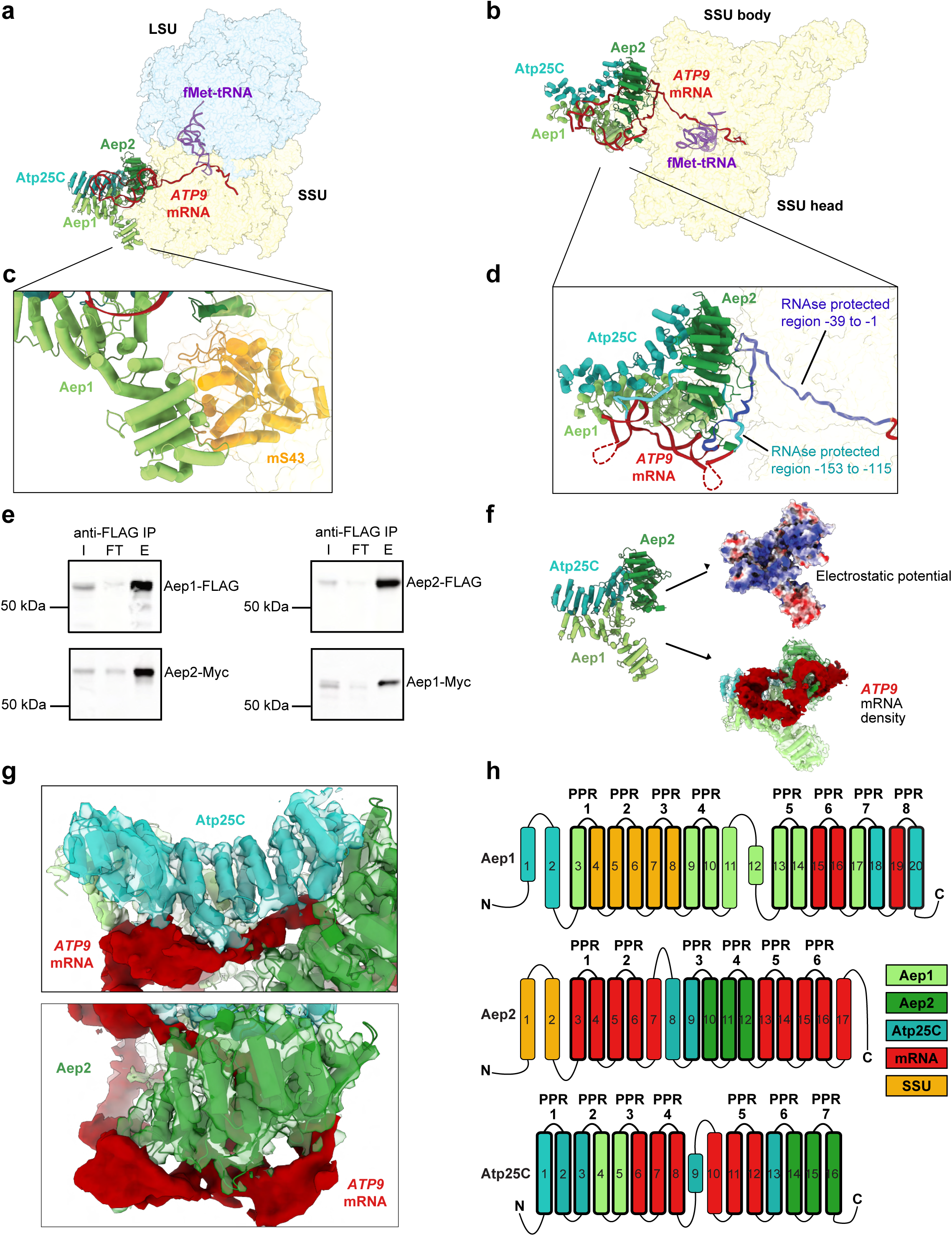
Interactions of the *ATP9* TAs with each other, the mitoribosome, and mRNA. **a,** Side view of *ATP9* TAs bound to the mitoribosome with transparent surfaces of the large subunit (LSU) and small subunit (SSU) colored in light blue and yellow, respectively. Aep1 (light green), Aep2 (green), and Atp25C (cyan) are shown bound to the SSU and *ATP9* mRNA (red). The initiator tRNA is colored in purple. **b,** Top view of *ATP9* TAs bound to the SSU. Color-coded as in a. **c,** Zoom in from panel a of interactions of Aep1 with the SSU protein Mrp1 (mS43) in yellow. **d,** Zoom in from panel b highlighting the interactions of the PPR domains of Aep1, Aep2, and Atp25C with the *ATP9* mRNA. The mRNA is color-coded with the regions predicted to correspond with the Aep1 and Aep2 sel-mitoRP footprints from Fig. 2a. Putative locations of predicted stem loop structures (Extended Data Fig. 8b) not resolved in the density map are denoted with a dashed line. **e,** Co-IP of Aep1 and Aep2 from whole cell lysate without crosslinking or RNase treatment. I= input, FT = flowthrough, and E = elution. **f,** Surface map of the *ATP9* TA complex structure highlighting the positively charged residues (blue) that interact with the mRNA surface depicted below. **g,** Zoom in of Atp25 (top) and Aep2 (bottom) PPR-RNA contact sites. **h,** Linear diagram highlighting the different PPR motifs that were resolved in the structure and the contacts that they make with either the mRNA or other proteins. Each alpha-helix is numbered and color-coded with its interacting partner.

Our structures revealed that Aep1 makes direct contacts with the mitoribosome through Mrp1 (mS43), a component of the SSU that is unique to the yeast mitoribosome and has been found to genetically interact with TAs^25^ (Fig. 5c). Aep2 and Atp25C make few direct contacts with the mitoribosome and are mostly linked to the mitoribosome through their interactions with Aep1. The PPR domains from each of the three proteins in the heterotrimer come together to form an RNA-binding scaffold. We were able to visualize mRNA density extending from the tRNA binding sites, where the initiator fMet-tRNA presumably base pairs with the AUG start codon, through the mRNA channel to the MCE, where the mRNA exits the mitoribosome and wraps around the TA-complex (Fig. 5a,b,d).

### TA-mitoribosome structures are consistent with their 5’ UTR footprints

The *ATP9* 5’ UTR footprints that we recovered in the Aep1 and Aep2 sel-mitoRP datasets can be explained by the structure of the *ATP9* TA-complex bound to the 5’ UTR (Figs. 1d and 2a). One set of RNA footprints, described above, abuts the start codon, and are consistent with an initiating mitoribosome bound to the TA complex. This footprint arises from the combined protection of the mitoribosome from the start codon until the MCE and the TA-complex, which protects an additional 20 nucleotides upstream (Figs. 2a and 5d). Sel-mitoRP of Aep1 and Aep2 also exhibited footprints approximately 130 nucleotides upstream of the start codon, which initially were unexpected (Figs. 1d and 2a). However, these footprints can be explained by two predicted stem loop structures between the UTR footprints (Extended Data Fig. 8b), bringing the upstream footprint much closer to the start codon in three-dimensional space (Fig. 5d). We did not visualize the loop portions of the stem loops in our structure, as the mRNA density was broken in the region where the loops would be found, probably due to flexibility of portions of RNA not bound by the TAs (Fig. 5d).

Binding of the 5’UTR of *ATP9* is achieved through a positively charged surface of the Aep1-Aep2-Atp25C complex that is complementary to the mRNA conformation (Fig. 5f). PPR motifs 6 and 8 of Aep1 make contact with the 5’ UTR of *ATP9*, while PPR motifs 1-3 of Aep1 do not bind the RNA, but instead establish protein-protein contacts with the mitoribosome (Fig. 5c,h). Furthermore, Atp25C makes extensive contacts with the 5’ UTR through PPR motifs 3, 4, and 5 (Fig. 5d,g,h). Atp25C thereby forms a bridge between Aep1 and Aep2 through its protein-protein and protein-RNA interactions relying on its PPR repeats (Fig. 5d,g,h). Aep2 localizes adjacent to the MCE and makes the most contacts with the 5’ UTR through PPR motifs 1, 2, 5, and 6, including an RNA loop positioned between Aep2 and the surface of the mitoribosome (Fig. 5d,g,h). This RNA loop likely contributes to the selectivity of Aep2 in recognizing and binding the *ATP9* mRNA to guide the mitoribosome to the correct start codon during initiation. Together, Aep1 and Aep2 sel-mitoRP and the *ATP9* TA mitoribosome structure demonstrate that the *ATP9* TAs link the *ATP9* 5’ UTR and the MCE to position the *ATP9* mRNA for initiation.

### *ATP9* TAs interact with the mitoribosomes during translation initiation and before elongation

Atp9 biogenesis is special, as it requires the assembly of ten identical subunits into a ring structure, which is the central part of the rotor component of ATP synthase. A possible scenario is that *ATP9* mRNA is translated by the same mitoribosome over and over to produce many copies of the protein to facilitate assembly. This in turn could be aided by a more constant binding of the *ATP9*-TA complex to the ribosome. To test this, we next asked whether the complex is directly released from the mitoribosome upon commencement of translation elongation, or whether it would stay bound longer. To address this question, we performed affinity purification of Aep2-bound mitoribosomes from an Aep2-3xFLAG-tagged strain actively respiring and translating (Fig. 6a and Extended Data Fig. 10). Cryo-EM analysis of these Aep2-mitoribosome particles yielded a prevailing class showing a highly similar density map as the *ATP9* TA-mitoribosome complex described above (Fig. 6b). Indeed, overlay of the TA-bound SSU from these two cryo-EM datasets revealed that the TA complex made nearly identical interactions with the mitoribosome (Fig. 6c,d), with a similar arrangement of the bound mRNA and tRNA bound to the P- or E-site. In addition, our overlapping structures obtained through orthogonal means indicate that the *ATP9* TAs bind only at early stages of translation to newly initiating mitoribosomes, consistent with sel-mitoRP data showing a rapid depletion of Aep1 and Aep2 from the mitoribosome after initiation (Figs. 1d and 6e).

**Figure 6:**
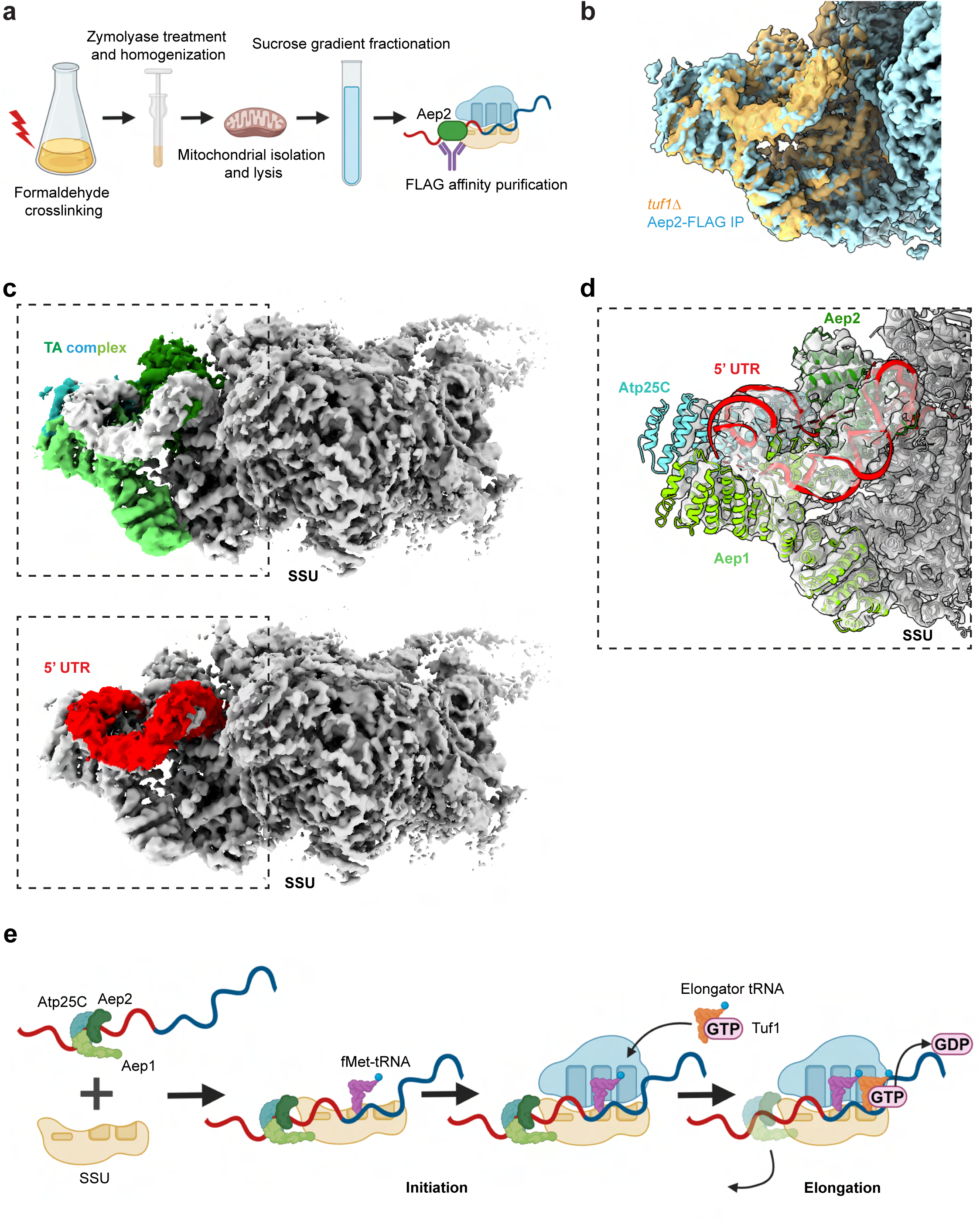
TA complexes preferentially engage with initiating mitoribosomes to position their target mRNA for translation initiation. **a,** Illustration representing different steps for the sample preparation of the affinity purified Aep2-bound mitoribosomes. **b,** Overlay of the cryo-EM density of *ATP9* TA complex obtained from the Aep2-affinity purified mitoribosomes with the *ATP9* TA complex bound mitoribosome obtained from *tuf1Δ* yeast cells. The affinity purified mitoribosomes are colored blue and the *tuf1Δ* mitoribosomes are colored yellow. **c,** Composite map of the SSU, obtained from the Aep2-affinity purified mitoribosomes, bound to *ATP9* TA complex highlighting the color coded Aep1-Aep2-Atp25C density (left top) and 5’UTR density in red (left bottom). **d,** The structure of the *ATP9* TA complex from Figure 3e fitted into the density of the Aep2-affinity purified mitoribosomes. **e,** Model of mRNA recruitment and positioning by the *ATP9* translational activator (TA) complex. The TA complex engages with the 5’ UTR of its target transcript and positions the start codon within the P-site of the SSU. The TA complex then departs the mitoribosome during early translation elongation.

## Discussion

The faithful choice of the start codon by ribosomes is key for correctly identifying and reading the intended open reading frame during protein synthesis. Here, we have identified how TAs support this key step during translation initiation in yeast mitochondria. The combination of sel-mitoRP analyses and cryo-EM structures demonstrate that TAs bind specifically to their target transcript and to the mitoribosomal small subunit to position the start codon in the P-site of the mitoribosome for translation initiation (Fig. 6e). The TAs bind to their target transcripts in the 5’ UTR approximately 30 nucleotides upstream of the start codon (Fig. 2). The precise position of the TA binding is likely aided by predicted RNA secondary structures around the binding sites and determines the distance between the TA binding site and the start codon (Extended Data Fig. 8). Binding of the TAs to the MCE then allows for the alignment of the mRNA into the mRNA channel for translation initiation. Thereby, mitochondrial TAs replace the molecular function of the Shine-Dalgarno mechanism, which is used by the ancestral bacterial system.

TAs were identified early during the genetic mapping of genes affecting mitochondrial functions and were later shown to play crucial roles for translation of individual mitochondrial-encoded mRNAs. Through their molecular function during initiation, they not only are necessary for mitochondrial translation, but also play key roles in organizing and regulating mitochondrial protein synthesis^6^. Data obtained over the years have suggested that TAs can also localize the synthesis of specific OXPHOS subunits to distinct sites in the inner membrane to presumably increase assembly efficiency^38,39^. Likewise, mitochondrial translation of OXPHOS proteins must be carefully coordinated with synthesis of nuclear-encoded OXPHOS subunits to ensure the stoichiometric assembly of OXPHOS complexes^2^. Recent work has demonstrated that *COB* translation is regulated by a feedback loop which depends on the influx of nuclear-encoded subunits and maintains the balance between nuclear- and mitochondrial-encoded subunits of Complex III^12,13^. Interestingly, Cbp1, which has been shown to be involved in this feedback loop, associated with mitoribosomes during *COB* translation initiation, but was not as selectively enriched on its target transcript as the other PPR TAs (Fig. 1c,d). This finding agrees with previous work showing that *COB* TAs can bind to the mitoribosome even in the absence of *COB* mRNA, arguing that *COB* TAs are bound to the mitoribosome even when not actively playing a role in translational activation^13^. Consistently, *COB* TAs engage with the mitoribosome through the large subunit (LSU) of the mitoribosome whereas our structures for the *ATP9* and *ATP8* TAs show that they bind to the SSU. Additionally, Cbp1 is found in a high molecular weight complex with Pet309, a *COX1* TA^40^. In agreement with these data we see a strong enrichment peak for Cbp1 in the 5’ UTR of *COX1* that overlaps with the Pet309 binding peak (Fig. 1c). The differences in how Cbp1 and potentially the other *COB* TAs engage with the mitoribosome argue that *COB* translation is regulated differently than other mitochondrial mRNAs.

Functional genetic studies have implied that TAs interact directly with the mitoribosome and the 5’ UTR of their target transcript, but biochemical and structural evidence of these interactions was lacking. The first cryo-EM structure of the yeast mitoribosome found an unresolved density at the extended MCE containing mitoribosomal-specific proteins^41^. It was proposed that this density was a heterogeneous mixture of TAs and that the MCE of the yeast mitoribosome could serve as a docking platform for TAs. Consistently, several TAs, including Aep1 and Pet111, were shown to populate the MCE by proximity labeling proteomics^9^. Our structural analysis of initiating mitoribosomes determined that the *ATP9* TAs (Aep1, Aep2, and Atp25C) form a heterotrimer that binds to the mitoribosomal protein Mrp1 at the MCE (Fig. 5c). The structure of Aep3 bound to an initiating mitoribosome revealed that it binds to the mitoribosome through Mrp51 (Fig. 4c). Hence, these data demonstrate that the expanded MCE of the yeast mitoribosome, which contains Mrp1 and Mrp51, serves as a docking site for the yeast TAs.

Our cryo-EM structures revealed how TAs engage with the mRNA during translation initiation by direct binding of the *ATP9* and *ATP8* 5’ UTRs by their respective TAs (Figs. 4a-f and 5a-h), which corresponded well with the footprints identified near the start codons of those transcripts (Fig. 2a,b). Indeed, we found that most of the TA-mitoribosome complexes exhibited footprints just upstream of the start codon that are likely due to the RNA protection afforded by TAs at their binding sites. Mitochondrial TAs must adapt to meet the needs of the divergent mitochondrial transcriptomes across species. Plant PPR proteins bind RNA in a sequence-specific manner dictated by the plant PPR code, whereby residues located in the loops between the two alpha helices in the PPR motif are used to decipher a specific nucleotide in an RNA sequence^42,43^. Sel-mitoRP data of this work indicate that most of the yeast PPR TAs bind their target mRNAs with high selectivity. Given that mitochondrial UTRs have a highly enriched A/U content, it is unlikely that PPR binding follows a strict sequence-specific RNA-binding mode. Indeed, the structures of TA-mitoribosome complexes with bound mRNA show that 5’ UTR binding by PPR proteins involves not only single strand interactions, but also the recognition of more complex secondary structures including double strands that presumably bring together annealing sequences spread on the RNA in three-dimensional space (Figs. 4c-f and 5d,f-h and Extended Data Fig. 8a,b). These interactions would be in line with the TA-mitoribosome footprints further upstream for *ATP9 and COX1* (Fig. 2a,c). Moreover, a T to C conversion at -16 in the *ATP9* 5’ UTR rescues a temperature sensitive allele of Aep2^44^ and is predicted to disrupt the putative stem loop that forms upstream of the start codon that would sterically prevent ribosome binding (Extended Data Fig. 8c). Of note, we found that PPR motifs of these TAs do not exclusively interact with RNA but are also involved in establishing protein-protein contacts, *e.g.* between the components of the *ATP9*-TA complex or to the mitoribosome (Figs. 4c and 5c). How RNA structure impacts TA binding and mitoribosome initiation represents an interesting future direction.

Evolution of the human mitochondrial genome has led to the complete loss of 5’ UTRs on protein-coding transcripts, such that if TAs exist for each human mitochondrial transcript they have changed so much in structure and function from their yeast counterparts as to be unrecognizable. TACO1 was initially assumed to be a TA of mammalian *COX1*^45^, however it was recently shown to act during translation elongation^46^. LRPPRC, a mammalian PPR protein, binds to the mitoribosome at the mRNA channel entrance and impacts translational efficiency of several mitochondrial transcripts^47^ by a yet poorly understood mechanism. Future work performing sel-mitoRP for LRPPRC and other putative translational regulators in human mitochondria will be critical to understanding how mitochondrial translation has adapted in response to the loss of the regulatory 5’ UTRs that are so critical for translation initiation in yeast. This approach will provide crucial insights into the evolutionary strategies employed to maintain efficient and specific mitochondrial translation across diverse eukaryotic lineages.

## Methods

### Yeast strain construction

Gene tagging and deletion was performed using standard transformation of PCR products^48^. All transformants were verified by colony PCR and western blotting. Epitope tags were introduced at the 3’ end of the endogenous locus using a scarless loopout method^49,50^.

For the strain with mitoribosomes stalled at initiation, a KanMX cassette was used to delete *TUF1* in a W303 strain background carrying the plasmids YEplac181-RNR1 (LEU2 selection) and pRS316-VAR1 (URA3 selection)^51^. All strains, plasmids, and primers used to generate the strains in this study are listed in Supplementary Tables 3, 4, and 5, respectively.

### Selective mitoribosome profiling

Yeast were grown in YPGal (1% yeast extract, 2% peptone, 2% galactose), pH 5.0, until an OD600 of approximately 1. 150 mL of culture was snap chilled prior to formaldehyde crosslinking by pouring the culture into a prechilled 500 mL centrifuge bottle filled with 50 grams of frozen, crushed ice cubes made from MilliQ water. The culture was quickly swirled in the centrifuge bottle and placed on ice. 10 mL of 16% formaldehyde was added to the chilled culture and the crosslinking proceeded for 1 hr on ice with swirling every 15 minutes. 11 mL of 2.5 M glycine was added to the culture to quench the crosslinking reaction. The culture was then pelleted and washed twice in 40 mL of ice cold crosslinking wash buffer (50 mM HEPES pH 7.5, 50 mM NH_4_Cl, 10 mM MgCl_2_). The pellet was then resuspended in 4 mL of crosslinking lysis buffer (50 mM HEPES pH 7.5, 50 mM NH_4_Cl, 10 mM MgCl_2_, 1.5X Complete EDTA-free protease inhibitor cocktail, 0.5% lauryl maltoside) and dripped into a 50 mL conical filled with liquid nitrogen to form frozen droplets.

The frozen cells with lysis buffer were then mechanically lysed using liquid nitrogen-chilled 50 mL canisters for six cycles of 3 min at 15 Hz using a Retsch MM301 mixer mill. The canisters were chilled in liquid nitrogen in between cycles to keep the cells frozen during the lysis. The frozen cell powder was transferred to a 50 mL conical and thawed in a room temperature water bath. The thawed cell lysate was diluted with 2 mL of fresh crosslinking lysis buffer.

To isolate ribosome protected RNA fragments, the lysates were treated with 300 Units of RNase I (Epicentre) for 30 minutes in a room temp water bath, swirling halfway to mix. 600 units of SUPERase IN RNase inhibitor was added to the lysate to stop RNase digestion. The RNase-treated lysate was then centrifuged at 3000 rpm for 5 min at 4 degrees C. The cleared lysate was then further clarified by transferring to 1.5 mL tubes and centrifuging for 15 min at 20,000 x G at 4 deg C. 3.5 mL of the clarified lysate was layered on top of 8 mL of sucrose cushion buffer (50 mM HEPES pH 7.5, 50 mM NH_4_Cl, 10 mM MgCl_2_, 1.5X Complete EDTA-free protease inhibitor cocktail (Roche), 24% sucrose). The lysate was then pelleted through the sucrose cushion to isolate ribosomes by centrifuging for 4.5 hours at 40,000 rpm at 4 deg C.

The sucrose cushion and cleared lysate was then removed and the ribosome pellet was resuspended in chilled mitoribosome wash buffer (50 mM HEPES pH 7.5, 50 mM NH_4_Cl, 10 mM MgCl_2,_ 0.1% Triton-X-100) overnight at 4 deg C with shaking.

The ribosome pellet was further resuspended by pipetting up and down several times and transferred to a 1.5 mL tube. The pellet was then incubated for 30 minutes at 4 deg C with end over end rotation to ensure that the pellet was well resuspended. The pellet was then centrifuged at 20,000 x G for 10 minutes to remove any insoluble residue. The clarified ribosome mixture was then added to 50 uL of packed Anti-FLAG M2 Affinity Gel (Millipore-Sigma) and incubated with end over end rotation for 3 hrs at 4 deg C. The resin was then washed 3 times with 1 mL of cold mitoribosome wash buffer. Following the last wash, the resin was resuspended with 1 mL of mitoribosome wash buffer using a cut pipette tip and transferred to a fresh 1.5 mL tube. The wash was removed and the resin was resuspended in 600 uL of mitoribosome wash buffer with 200 ug/mL 3x-FLAG peptide (Millipore-Sigma). The resin was then incubated for 1 hr at 4 deg C with rotation. To recover the elution, the resin slurry was passed over a Co-Star SpinX column for 1 min at 16,000 X G. The elution was then transferred to a fresh 1.5 mL tube.

To reverse the formaldehyde crosslinking, 30 uL of 20% SDS, 32 uL of 100 mM DTT, and 14 uL of 0.5 M EDTA were added to the elution, which was then heated for 45 min at 70 deg C. The RNA footprints were extracted following reversal of the crosslinking using an equal volume of acid-phenol chloroform. Ribosome footprints were then isolated and a cDNA library was prepared for Illumina short read sequencing as previously described ^52^.

### Selective ribosome profiling data analysis

Reads were trimmed to remove ligated 3’ linker (CTGTAGGCACCATCAAT) with Cutadapt^53^. Reads without linker were discarded and the unique molecular identifier (UMI, consisting of 4 random nucleotides introduced at the 5’ end with the RT primer and 10 random nucleotides introduced at the 3’ end with the linker) was then extracted from remaining reads using a custom script. For all Aep1 and Aep2 samples, synthetic RNA oligos of 40 and 33 nucleotides (UAACAACAUUCAUUAUGAAUGAUGUACCAACACCUUAUGC and UAACAACAUUCAUUAUGAAUGAUGUACCAACAC) were used as size markers in a separate gel lane during mitoribosome footprint size selection. Reads arising from migrating synthetic oligos were removed *in silico* using BBTools’ BBDuk (sourceforge.net/projects/bbmap/) with a k-mer length of 30. Reads mapping to abundant non-coding RNAs (rRNA, tRNA) were filtered out after alignment using bowtie1^54^. Remaining reads were aligned to the *S. cerevisiae* genome assembly R64 (UCSC: sacCer3) using STAR 2.7.3^55^. PCR duplicates were identified by their UMI and removed using a custom script. Ribosome A site positions were determined using an offset from the 3’ end of each read, depending on its length: 37:[-15], 38:[-16], 39:[-17], 40:[-17], 41:[-17]. Shorter read lengths have ambiguous offsets and were omitted.

Unless otherwise noted, read counts at each A site position were summed across replicates (2 to 6 replicates for each sample) and normalized by total mitochondrial mRNA-mapping reads (rpm). Enrichment over total mitoribosomes was then calculated by summing rpm values in a sliding 9-nt window and dividing by the corresponding sum in the total (Mrps17) dataset. A threshold was set so that if more than 3 positions had no coverage in either sample the enrichment value was set to 0. This was to avoid spurious signal due to low coverage. GC content was calculated for a sliding 6-nt window.

### Western blotting

5x SDS-PAGE loading buffer was added to samples at a final concentration of 1x and samples were boiled for 5 min at 95 deg C prior to loading onto NuPAGE Bis-Tris gels (Invitrogen). Protein transfer to a nitrocellulose membrane was carried out using a Trans-Blot Turbo (Bio-Rad), and the membranes were blocked in 5% milk/TBST (0.1% Tween-20). 3x-FLAG-tagged proteins were detected with mouse anti-FLAG (Millipore-Sigma, F1804, 1:1000), 3x-HA tagged proteins were detected using mouse anti-HA (Invitrogen, 26183, 1:1000), and 3x-Myc tagged proteins were detected using mouse anti-Myc (Millipore-Sigma, 05-724, 1:500). IRDye 800CW goat anti-mouse secondary antibodies (LICORbio, 925-32210, 1:5000) were used to image protein bands on a LICOR machine.

### Mitochondrial isolation and preparation of *tuf1Δ* mitoribosomes

Mitochondria were isolated according to a previously established protocol^56^. In short, yeast was cultured in 8 L of synthetic media containing 1.7 g/L yeast nitrogen base, 5 g/L (NH_4_)_2_SO_4_, 20 μg/ml of adenine, uracil and arginine, 15 μg/ml of histidine and 30 μg/ml lysine supplemented with 2% glucose. Cells were grown overnight at 30°C and 170 rpm to an OD_600_ of 3 before harvesting at 3000x*g* for 5 min, washed with distilled water and resuspended in MP1 buffer (2 ml/g wet weight of 0.1 M Tris-base and 10 mM dithiothreitol). After incubation for 10 min at 30°C, cells were washed with 1.2 M sorbitol, resuspended in MP2 buffer (6.7 ml/g wet weight of 20 mM KPi pH 7.4, 0.6 M sorbitol and 3 mg/g wet weight of zymolyase (Seikagaku Biobusiness, Tokyo, Japan) and incubated at 30°C for 1 h. Cells were harvested at 3000x*g* for 5 min at 4°C, resuspended in MP3 buffer (13.4 ml/g wet weight of 0.6 M sorbitol, 10 mM Tris pH 7.4, 1 mM EDTA and 1 mM PMSF) and homogenized with 2 x 10 strokes using a tight-fitting homogenizer (Sartorius Stedim Biotech S.A., France). The homogenate was centrifuged two times at 3000x*g* for 5 min at 4°C before mitochondria were harvested by centrifugation at 15.000x*g* for 15 min at 4°C. The mitochondrial pellet was resuspended in SH buffer and frozen in liquid nitrogen before storage at -80°C.

For isolation of mitochondrial ribosomes, approximately 50 mg of mitochondria were thawed on ice and centrifuged at 10.000x*g* for 10 min at 4°C before resuspension in lysis buffer (25 mM HEPES pH 7.4, 100 mM KCl, 20 mM MgOAc, 1% DDM, 1 mM PMSF and 1x Complete protease inhibitor (Roche) in RNAse free H_2_O) and incubation on ice for 10 min. The lysate was clarified by centrifugation two times at 20.000x*g* for 10 min 4°C before being loaded on a sucrose cushion (1.2 M sucrose, 25 mM HEPES pH 7.4, 100 mM KCl, 20 mM MgOAc, 0.1% DDM and 2 mM DTT in RNAse free H_2_O) and centrifuged in a TLA 120.2 rotor at 75.000 rpm (245.000 g) for 3 h at 4°C. The mitoribosome pellet was resuspended in a grid buffer (25 mM HEPES pH 7.4, 100 mM KCl, 20 mM MgOAc and 0.02% DDM in RNAse free H_2_O) and clarified through centrifugation at 20.000x*g* for 10 min at 4°C. Sample was diluted to an OD_260_ of 10 corresponding to approximately 230 nM.

### Data acquisition of *tuf1Δ* sample

3.5 μl of sample was applied to glow-discharged (20 mA for 60s) QuantiFoil R2/2 Cu 300 mesh grids pre-coated with 3 nm carbon, using 30 s wait time and 3 s blot time at 100% humidity and 4°C, before plunge freezing in liquid ethane using a Vitrobot Mark IV (FEI, Thermo Fisher Scientific). Two data sets containing 26,435 and 25,922 movies were collected on a FEI Titan-Krios G3 (Thermo Fisher Scientific) transmission electron microscope operating at 300 keV and equipped with a Gatan K3 direct electron detector and a GIF quantum energy filter with a slit width of 20 eV. The movies were acquired using a pixel size of 0.828 Å, and a nominal magnification of 165.000x, using the software EPU, with 40 frames per movie, a defocus range of 0.4-2.6 μm and a total electron dose of 38 eÅ^-2^.

### Image processing of *tuf1Δ* sample

Image processing was performed using *cryoSPARC* v4.3.1 (Structura Bio) (Extended Data Fig. 6) with initial pre-processing of the two data sets following a standard processing pipeline containing Patch motion correction, Patch CTF estimation and curation of exposures to remove micrographs of low quality and bad CTF estimations. Particles were picked using reference-free blob picking which was followed by particle extraction with a box size of 600 Å which was binned 2x (1.64 Å per pixel). Several rounds of 2D classification were performed to generate good classes for template-based particle picking. Further rounds of 2D classification were followed by *ab-initio* 3D reconstruction and heterogenous refinement. Classes containing unaligned particles, cytosolic ribosomes and LSU only were discarded and mitoribosome monosome particles from the two datasets were merged at this stage. The merged monosome particles were subjected to another round of 2D classification, *ab-initio* 3D reconstruction and heterogenous refinement that resulted in a stack of 461,825 particles. This pool of particles was first subjected to homogenous refinement followed by focused unaligned 3D classification using a mask on the mitoribosome subunit interface to look for tRNA occupancy. No class with tRNA density in the A-site was observed, which served as a good validation that this pool of mitoribosome particles were not elongating and likely stalled at initiation. Further, the 3D density acquired from the homogenous refinement showed a weak additional density at the mRNA exit channel on the SSU. The particles were therefore subjected to homogenous refinement with a binary mask covering the SSU and area around the mRNA exit channel. This was followed by unaligned 3D classification using 20 classes and a focused mask around the additional density. This resulted in two classes with a visibly similar larger density, which was subsequently merged, and one class with a smaller density all located at the mRNA exit channel. The two distinct classes of particles were used for another round of unaligned 3D classification, followed by re-extraction to a box size of 640 Å (0.82 Å per pixel), as well as global and local CTF refinement. Finally, the two classes consisting of 24,558 and 22,558 particles were subjected to local refinement, using focused masks and a FSC cut-off of 0.143, which resulted in density maps with estimated resolutions of 3.8 Å and 3.7 Å, respectively.

### Model building and refinement

Using *AlphaFold2*^57^, we predicted the structures of yeast translational activator (TA) proteins, known to interact with the mRNA exit channel, to systematically test if they could be fitted into the additional densities found in our maps. For the larger additional density, we found that three TAs of the *ATP9* mRNA, namely Aep1, Aep2 and Atp25C, fit very well when comparing the overall architecture and conformations of α-helices. The initial model for each protein was first rigid-body fitted into the density map using ChimeraX v1.6 (UCSF) and subsequently real space refined using *PHENIX* v1.21-5207^58^. Each protein subunit was then visually inspected and manually adjusted according to their densities using torsion, planar peptide, trans peptide and Ramachandran restraints in *Coot v0.9.8.1*^59^. For the yeast mitochondrial ribosome, previously published structures of the SSU (PDB ID: 8OM4) and LSU (PDB ID: 5MRC) were used as initial models that were rigid-body fitted into the density maps and each protein and RNA subunit manually adjusted in *Coot* and real space refined using *PHENIX*. An extra density, corresponding to the *ATP9* mRNA, was visibly bound to the TA protein complex and extended into the decoding centre where it interacted with a tRNA. An RNA backbone consisting of polyU (U-A in double-stranded regions) was modelled in *Coot and* real space refined using secondary structure restraints in *PHENIX* into this fractured density for visualization purposes. A tRNA structure (from PDB ID: 5MRC) in the E-site was used as an initial model and its nucleotides were modified in *Coot* according to the nucleotide sequence of the yeast initiator fMet-tRNA. The full structure containing the mitoribosomal LSU and SSU, fMet-tRNA, mRNA and an Aep1-Aep2-Atp25C complex was subsequently real-space refined in *PHENIX* using a composite map of focused and locally refined maps

The second class visibly contained a single protein density, with PPR motifs, bound at the MCE in a similar position to Aep2. We found that the predicted structure for the *ATP8* translational activator Aep3 fit very well into this map, which also contained partial density for double- and single-stranded mRNA extending into the ribosome. A model containing Aep3 and mRNA bound to the mitoribosome was subsequently rigid-body fitted, manually adjusted and refined in the same way as the *ATP9* TA complex. Two regions predicted to consist of long flexible loops ranging between residues 33-52 and 531-558 were removed due to a lack of clear density in the map.

### Isolation of Aep2-ribosome complexes for cryo-EM

Six liters of yeast culture in YPGal pH 5.0 was grown at 30 deg C until an OD600 between 1 and 1.5 was reached. The six liters of culture was aliquoted in 2000 mL in 3 4-L flasks. 650 grams of crushed MilliQ Ice cubes were added to each flask. The flasks were quickly swirled and placed on ice. 58 mL of 37% formaldehyde was added to each snap-chilled culture. Crosslinking proceeded for 1 hr on ice and the flasks were swirled every 15 minutes to mix. 142 mL of 2.5 M glycine was added to each culture to quench the crosslinking. The culture was pelleted by centrifugation for 10 min at 3000 rpm at 4 degrees Celsius. The pellet was then washed in MilliQ water twice and weighed.

The cell pellet was then resuspended with prewarmed DTT buffer (100 mM Trizma Base, 10 mM DTT) at 2 mL/g of wet weight. The resuspended cells were then transferred to a 250-mL flask and incubated at 30 degrees Celsius for 20 minutes with 80 rpm shaking. The cells were pelleted at 3000xG for 5 minutes and resuspended in 7mL/g of Zymolyase buffer (1.2 M sorbitol, 20 mM potassium phosphate, pH 7.4) containing 3 mg of Zymolyase 20T (MP Biomedicals) per gram of wet weight. The resuspended cells were then transferred to a flask and shaken at 80 rpm at 30 degrees Celsius for one hour. The cells were then pelleted for 5 min at 3000xG and washed with fresh Zymolyase buffer without Zymolyase 20T. The cells were then resuspended in 6.5 mL/g of ice cold homogenization buffer (0.6 M Sorbitol, 10 mM Tris-HCl pH 7.4, 1X Protease Inhibitor Cocktail EDTA-free) and homogenized using a glass dounce homogenizer for 20 strokes on ice. Following homogenization, dilute the lysate with an equal volume of cold homogenization buffer. Cell debris and nuclei were then pelleted by spinning the lysate at 1500xG for 5 min at 4 degrees Celsius. The supernatant was then further clarified by an additional spin at 3000xG for 10 minutes. Using a SW41-Ti rotor mitochondria were pelleted from the supernatant at 12,000xG for 15 minutes. The mitochondrial pellet was then resuspended in 5 mL of lysis buffer without lauryl maltoside (50 mM HEPES pH 7.5, 50 mM NH4Cl, 10 mM MgCl2, 1.5X Protease Inhibitor Cocktail EDTA-free, 1% lauryl maltoside). The resuspended mitochondria were then flash-frozen in liquid nitrogen and stored at -80 degrees Celsius.

Six 10-50% sucrose gradients were prepared using 10% (50 mM HEPES pH 7.5, 50 mM NH4Cl, 10 mM MgCl2, 1X Protease Inhibitor Cocktail EDTA-free, 10% sucrose) and 50% (50 mM HEPES pH 7.5, 50 mM NH4Cl, 10 mM MgCl2, 1X Complete Protease Inhibitor Cocktail EDTA-free (Roche), 50% sucrose) sucrose solutions using the Biocomp Gradient Master using the 10-50% SW41 14s preset program. The isolated mitochondria were thawed in a room temperature water bath, inverted to mix, and placed on ice. 25 uL of Superase-In RNase Inhibitor and 1 mL of 5% lauryl maltoside were added to the isolated mitochondria. The mitochondria were then pipetted up and down with a 5 mL pipette tip and incubated on ice for 1 hour in order to lyse the mitochondria. The mitochondrial lysate was then clarified by centrifuging for 20 minutes at 20,000xG at 4 degrees Celsius. 750 uL of clarified lysate was added slowly to the top of each 10-50% sucrose gradient. The gradients were centrifuged for 3 hrs at 40,000 rpm at 4 degrees Celsius in an SW41-TI rotor. The gradients were fractionated into 13 fractions of 850 uL and a Triax flow cell was used to measure the OD260 for RNA abundance within each fraction. The monosome fractions from each sucrose gradient were pooled and 18 uL of 5% lauryl maltoside, 50 uL of Superase-In RNase Inhibitor, and 210 uL of 50X Complete Protease Inhibitor Cocktail EDTA-free were added to the pooled fractions.

The pooled monosome fractions were then incubated overnight at 4 degrees Celsius with end-over-end rotation with 50 uL of packed anti-FLAG M2 affinity resin (Millipore-Sigma). The anti-FLAG affinity resin was then washed 3 times with 1 mL of mitoribosome wash buffer with lauryl maltoside (50 mM HEPES pH 7.5, 50 mM NH4Cl, 10 mM MgCl2, 0.1% lauryl maltoside). After the last wash, the FLAG resin was resuspended with 1 mL of elution buffer (50 mM HEPES pH 7.5, 50 mM NH4Cl, 10 mM MgCl2, 0.02% lauryl maltoside) and transferred to a clean 1.5 mL tube. The Aep2-FLAG ribosome complexes were then eluted from the resin with 600 uL of elution buffer with 200 ug/mL 3X-FLAG peptide for 1 hr at 4 degrees Celsius with end-over-end rotation. A second elution was carried out with 600 uL of elution buffer with FLAG peptide and the two elutions were pooled together and concentrated using a 500 uL 100K PES protein concentrator (Pierce).

### Data acquisition of Aep2-bound mitoribosome sample

For the Aep2-bound mitoribosomes, 3.5μl of the affinity purified mitoribosome sample was applied to glow-discharged R2/2 copper 400 mesh grids (Quantifoil) covered with a 5 nm layer of continuous carbon and plunge frozen in liquid ethane using a Vitrobot Mark IV (Thermo Fisher Scientific) that was set at 4 °C and 100% humidity with a 10 second wait time, 3 second blot time, and +6 blot force. The grids were first screened using a 200 kV Talos Arctica (Thermo Fisher Scientific). The final dataset was collected using a 300 kV Titan Krios (Thermo Fisher Scientific), equipped with a Falcon4i camera (Thermo Fisher Scientific) and Selectris energy filter of 10eV slit width, in counting mode at a nominal magnification of 105,000 corresponding to a calibrated pixel size of 1.19Å. A total of 15,728 EER movie frames were acquired with a total exposure time of 6s, corresponding to a total dose of 48.5 electrons/Å2. The defocus values were set from −0.8 to −2.0μm. Semi-automated data collection was performed with EPU (Thermo Fisher Scientific).

### Cryo-EM data processing of Aep2-bound mitoribosome

Data processing of the Aep2 bound mitoribosomes (Extended Data Fig. 10) was performed using both RELION v.4.0.1^60^ and with cryoSPARC v.4.3.1^61^. The movie frames were motion corrected using RELION’s own implementation and CTF estimation was performed using CTF Find 4.1^62^. A total of 874,238 particles were picked using automated particle picking in RELION v.4.0.1and extracted with a particle box size of 420 pixels, rescaled to 120 pixel with rescaled pixel size of 2.38 Å/pixel. Multiple rounds of 2D classification were performed to get rid of non-ribosomal particles. The selected particles from the best 2D classes were used for 3D classification and subsequent 3D refinement. The duplicated particles were removed from the refined particles, particles were reextracted to full pixel size of 1.19 Å/pixel and imported to cryoSPARC v.4.3.1 for further processing.

The imported 136,281 particles were subjected to homogeneous refinement in cryoSPARC v.4.3.1 and they were further taken for particle signal subtraction using masks generated by using either SSU or LSU from prior processing steps from RELION v.4.0.1. For both LSU and SSU, the signal subtracted particles were subjected to 3D classification with no alignments and the best classes were further subjected to homogeneous refinement followed by global and local CTF refinements resulting in density maps with estimated resolution of 2.9 Å for LSU and 3.6 Å for SSU respectively (FSC cut-off 0.143). To further obtain a better TA (Aep1-Aep2-Atp25 complex) density, particles from the refined SSU were taken and signal subtracted using a mask made to keep only the TA density. After the signal subtraction, 3D classification with no alignment followed by local refinement was performed resulting in density map with estimated resolution of 4.4 Å (FSC cut-off 0.143).

### Co-immunoprecipitation of Aep1 and Aep2 from whole cell lysate

Yeast were grown to an OD_600_ between 0.6 and 0.8 and harvested by rapid filtration and flash freezing in liquid nitrogen. The frozen cell pellet was then mixed with 4 mL of frozen crosslinking lysis buffer and mechanical lysis of the cells was performed as described for the sel-mitoRP. The frozen lysate was thawed at room temp and precleared by centrifugation at 3000 rpm for 5 minutes, followed by another preclearing step of 20,000xG for 15 minutes. Each sample was incubated overnight with 40 uL of packed anti-FLAG agarose resin with end-over-end rotation at 4 degrees Celsius. The flowthrough was removed and the beads were washed three times with 1 mL of mitoribosome wash buffer. The proteins were eluted with 200 ug/mL 3x-FLAG peptide for 1 hr at 4 degrees Celsius.

## Acknowledgements

We thank A. Amunts and Y. Zgadzay for help and advice on attempts to obtain the Aep2-mitoribosome structure. We thank C. Nuessmeier and G. Prakash for careful review of the manuscript. We thank T. Fox and members of the Churchman and Winston labs for advice and discussions. We acknowledge Harvard Center for Cryo-Electron Microscopy (HC2EM) for cryo-EM data collection and SBGrid for support of data processing. The cryo-EM data of *tuf1*Δ mitorobosome complexes were collected at the Swedish national cryo-EM facility funded by the Knut and Alice Wallenberg Foundation, Erling Persson and Kempe Foundations. This work was supported by National Institutes of Health grant R01-GM123002 (L.S.C.), the Swedish Research Council 2018-03694 and 2021-05545 (M.O.), the Knut and Alice Wallenberg foundation 2017.009 and 2019.0319 (M.O.), and the Packard Foundation (S.S), S.S. is an investigator with the Howard Hughes Medical Institute.

## Contributions

J.B.B. generated FLAG-tagged yeast strains for sel-mitoRP and performed the sel-mitoRP. J.B.B. and M.T.C. analyzed the sel-mitoRP sequencing results. B.W. performed the co-IP of Aep1 and Aep2 from whole cell lysate. J.B.B. purified the Aep2-bound mitoribosomes for cryo-EM. J.G. and D.S. made the cryo-EM grids for the Aep2 affinity purified mitoribosomes, collected the data, and determined the structure. A.C. generated the *tuf1*D strain, purified TA-mitoribosome complexes from this strain, made cryo-EM grids and performed cryo-EM screening, data collection, structure determination and model building with the help of U.R.. J.B.B., A.C. and D.S. prepared figures and a first draft of the manuscript. S.S., M.O. and L.S.C. supervised the project, interpreted the data and wrote the paper. All authors contributed toward the final version of the paper.

---

**Extended Data Figure 1:**
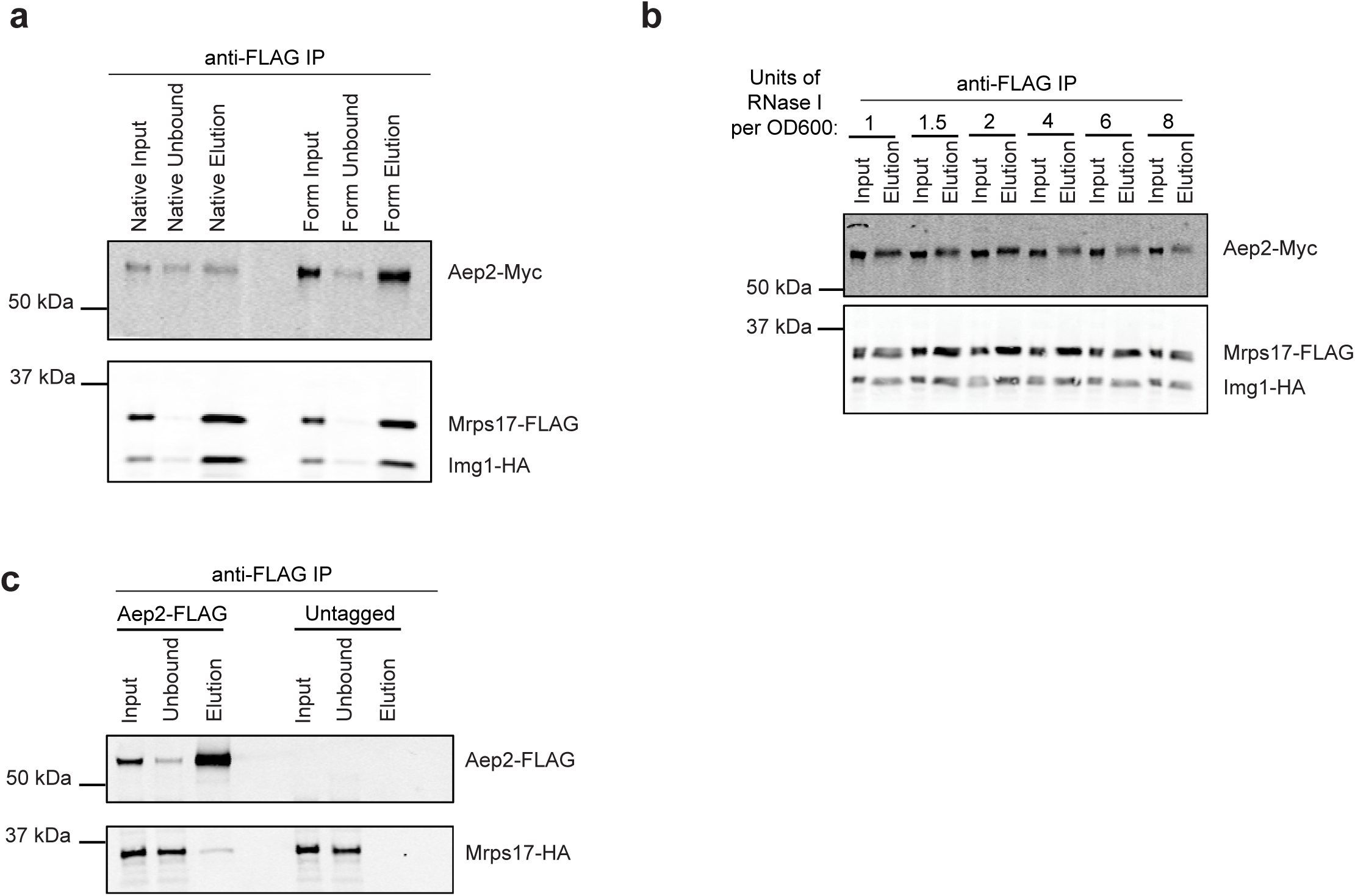
Optimization of TA co-purification with the mitoribosome. **a,** Western blot of Myc-tagged Aep2 co-purified with a 3X-FLAG-tagged mitoribosome from the monosome fraction of a 10-50% continuous sucrose gradient (Input). The co-IP was performed from yeast cells treated with or without formaldehyde crosslinking. **b,** Crosslinked yeast lysates were pretreated with increasing amounts of RNase I prior to affinity purification of Mrps17-FLAG mitoribosomes from the monosome fraction of a 10-50% sucrose gradient (Input). Continued interaction of Aep2-Myc with the mitoribosome was tested at each indicated concentration of RNase I. **c,** FLAG affinity purification from the monosome fraction of a 10-50% sucrose gradient was carried out for an Aep2-FLAG strain and untagged control demonstrating specific co-purification of the mitoribosome only when Aep2 is tagged.

**Extended Data Figure 2:**
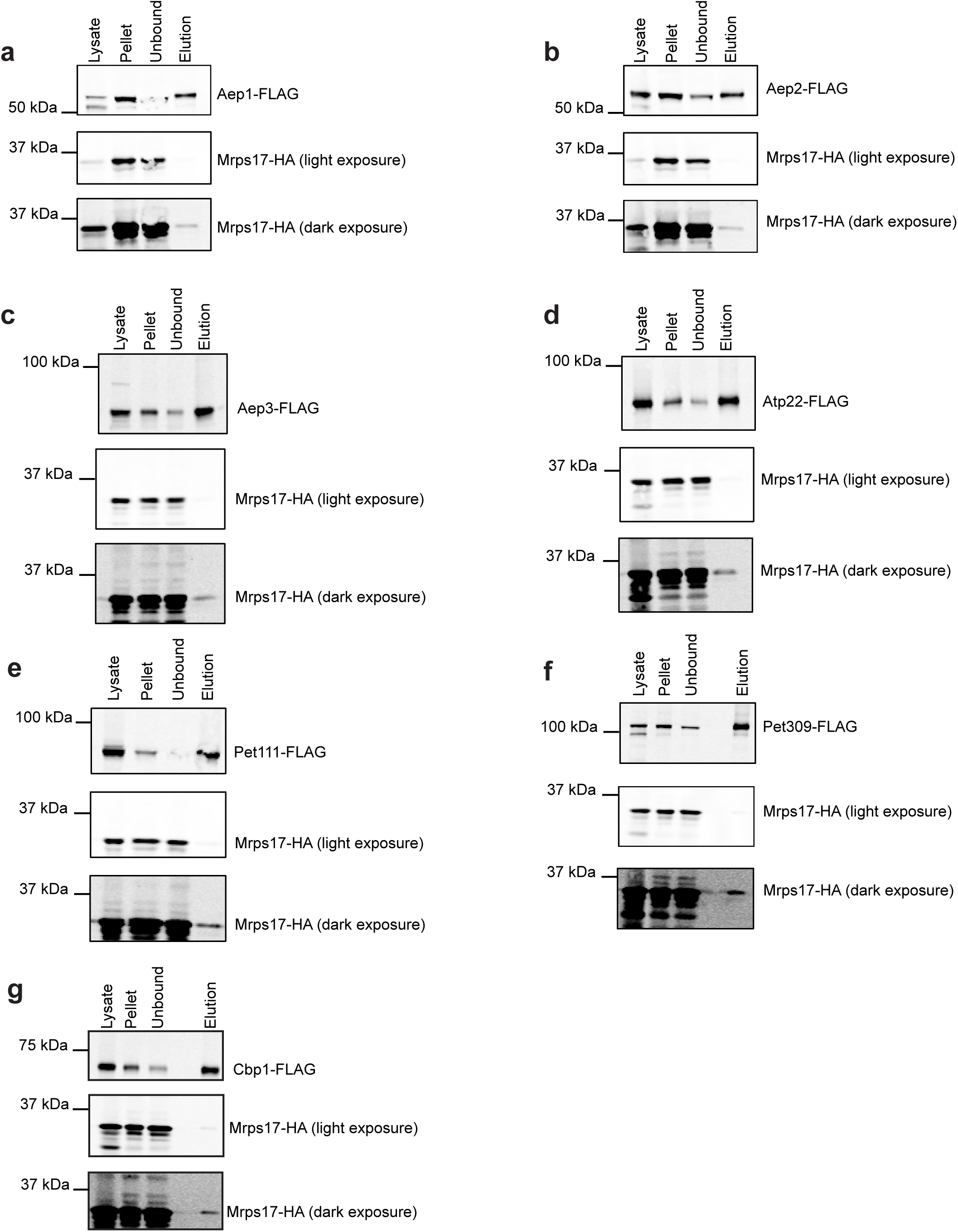
Co-IP of the mitoribosome with the purified Tas a-g, Western blots from the sel-mitoRP co-IP of the given FLAG-tagged TA and the HA-tagged mitoribosome. The blots were imaged on a LICOR machine and low contrast (low exposure) and high contrast (high exposure) images are shown. Lysate refers to the RNase treated lysate prior to its pelleting through a sucrose cushion. Pellet refers to the protein material recovered from the resuspended ribosome pellet. Unbound is the protein that was not bound to the anti-FLAG agarose resin. Elution is the material that was eluted off the anti-FLAG resin with 3X-FLAG peptide.

**Extended Data Figure 3:**
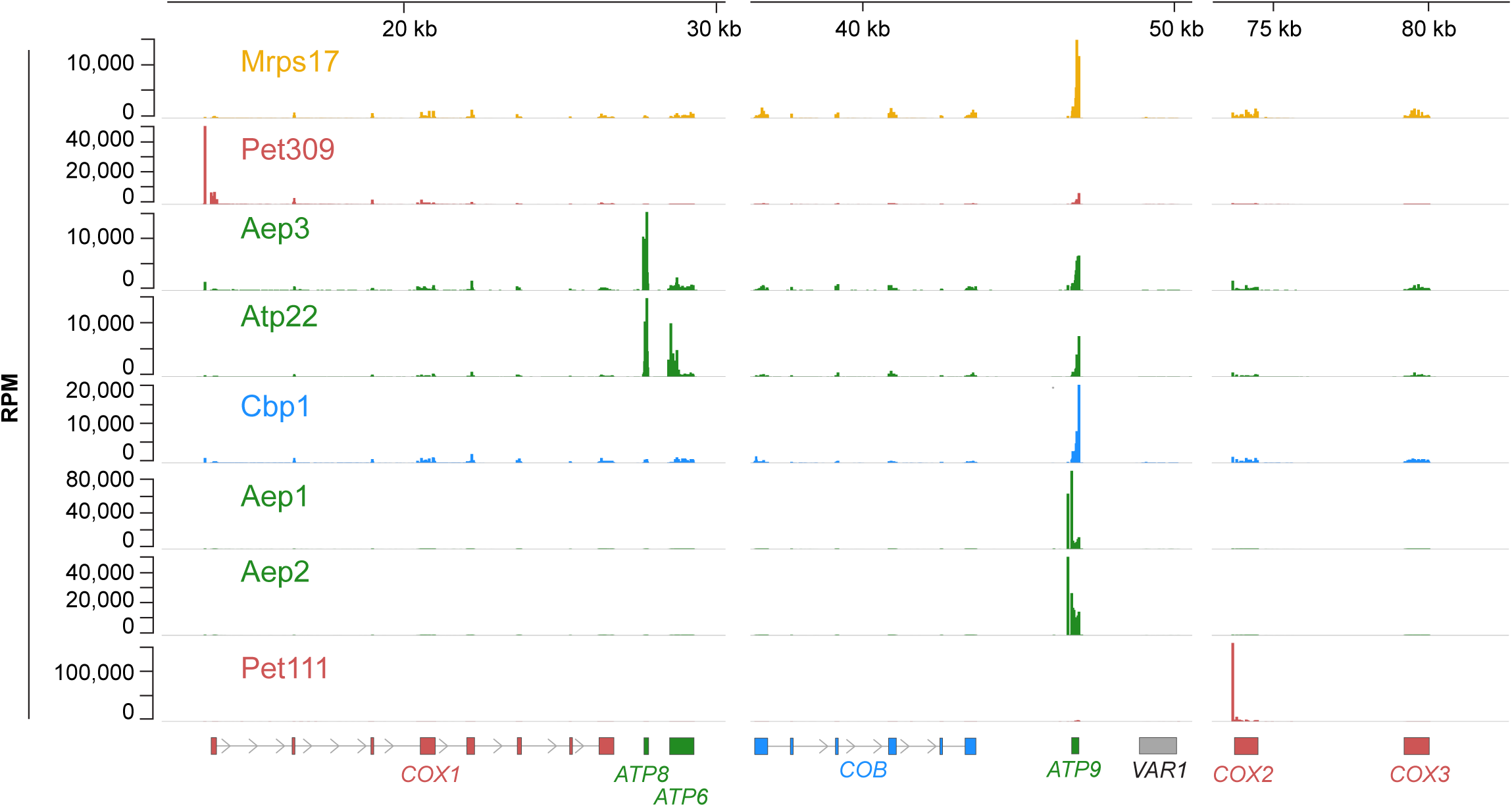
Sel-mitoRP data without Mrps17 normalization. Inferred A-sites for Mrps17 and all sel-mitoRP datasets normalized to total reads mapping to the mitochondrial genome (RPM). RPM = reads per million mapped reads

**Extended Data Figure 4:**
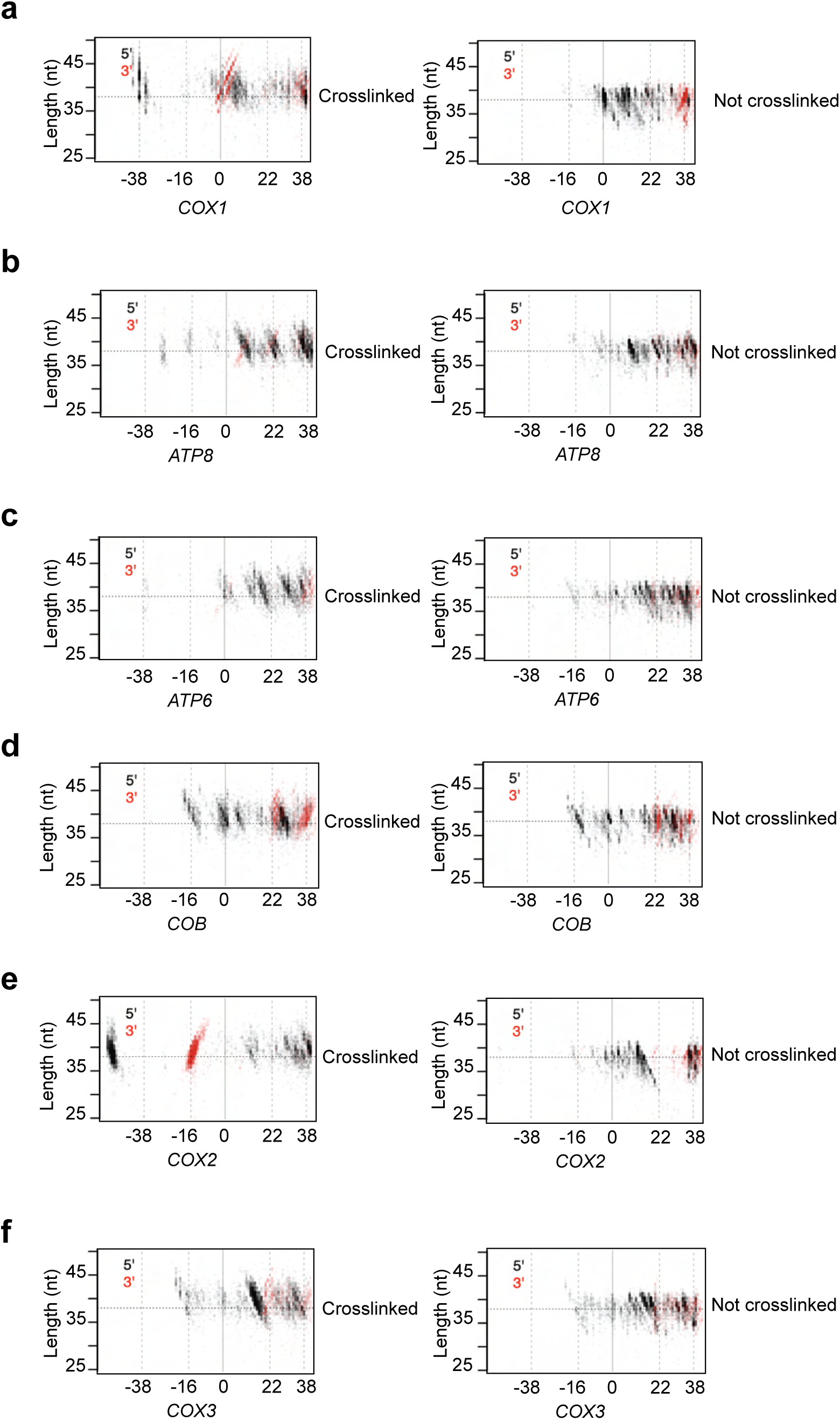
Crosslinking extends the 5’ end protection of initiating mitoribosomes a-f, Plots comparing total mitoribosome profiling either with or without crosslinking for the given transcript. 5’ ends of RNA footprints are plotted in black and 3’ ends are plotted in red. The horizontal dashed line indicates a standard initiating mitoribosome footprint size of 38 nucleotides. For all transcripts except for *COB* and *COX3*, a clear extension to the 5’ end of the initiating mitoribosome can be visualized upon the addition of formaldehyde crosslinking.

**Extended Data Figure 5:**
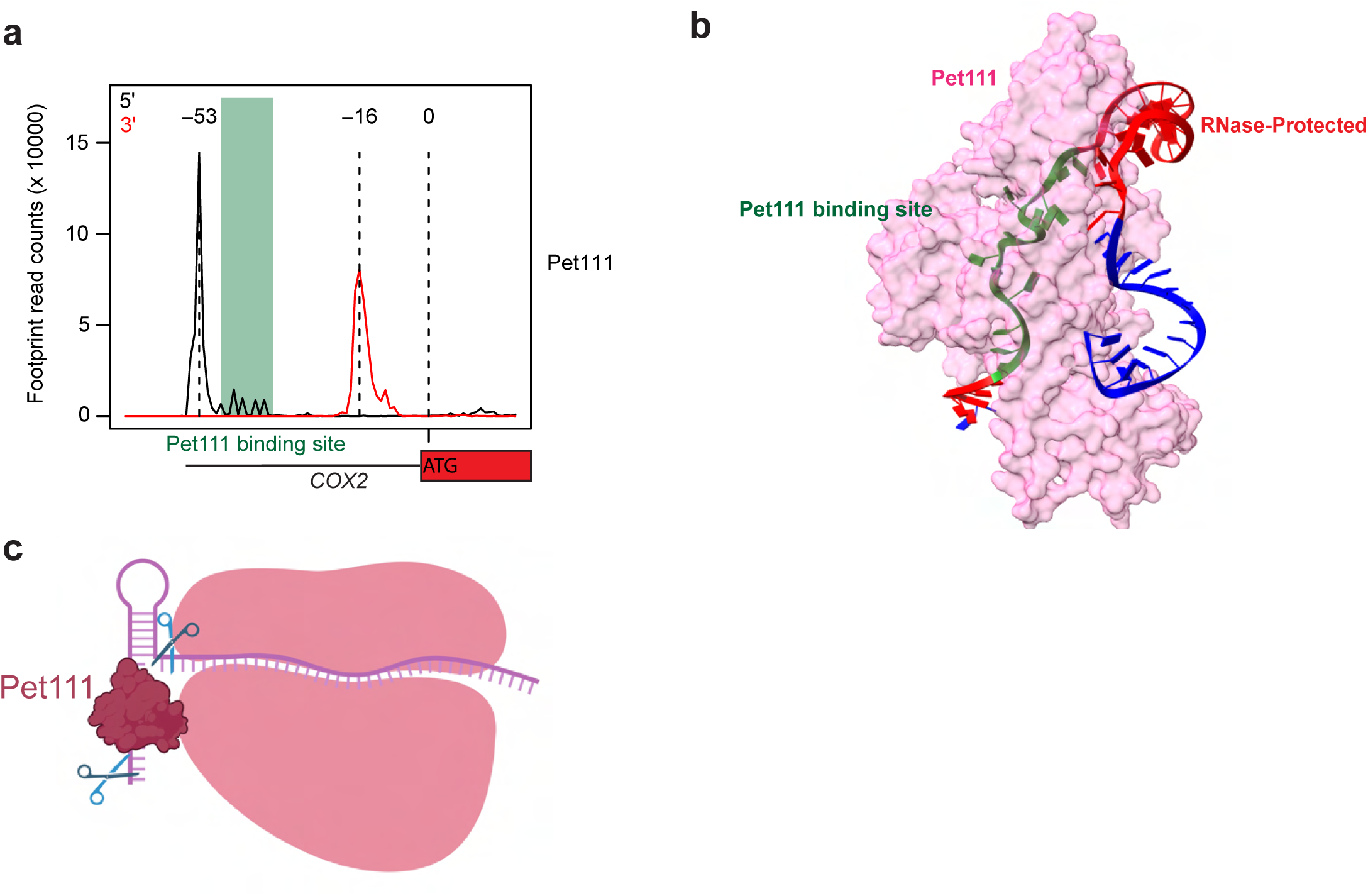
Structure prediction of Pet111 bound to the *COX2* 5’ UTR. **a,** The read counts for the 5’ and 3’ ends of Pet111 sel-mitoRP footprints are plotted in relation to the Pet111 binding site in the 5’ UTR of *COX2*. 5’ ends are plotted in black and 3’ ends are plotted in red. **b,** The AlphaFold3 structural prediction of Pet111 (magenta) bound to its target site in the *COX2* 5’ UTR. The Pet111 binding site is highlighted in green and the RNase-protected region (red) indicates the bounds of the 5’ and 3’ end peaks that surround the Pet111 binding site in panel a. **c,** Model of Pet111 bound to *COX2* upstream of the predicted stem loop structure with RNase I (scissors) cutting the RNA upstream of Pet111 and downstream of the stem loop.

**Extended Data Figure 6:**
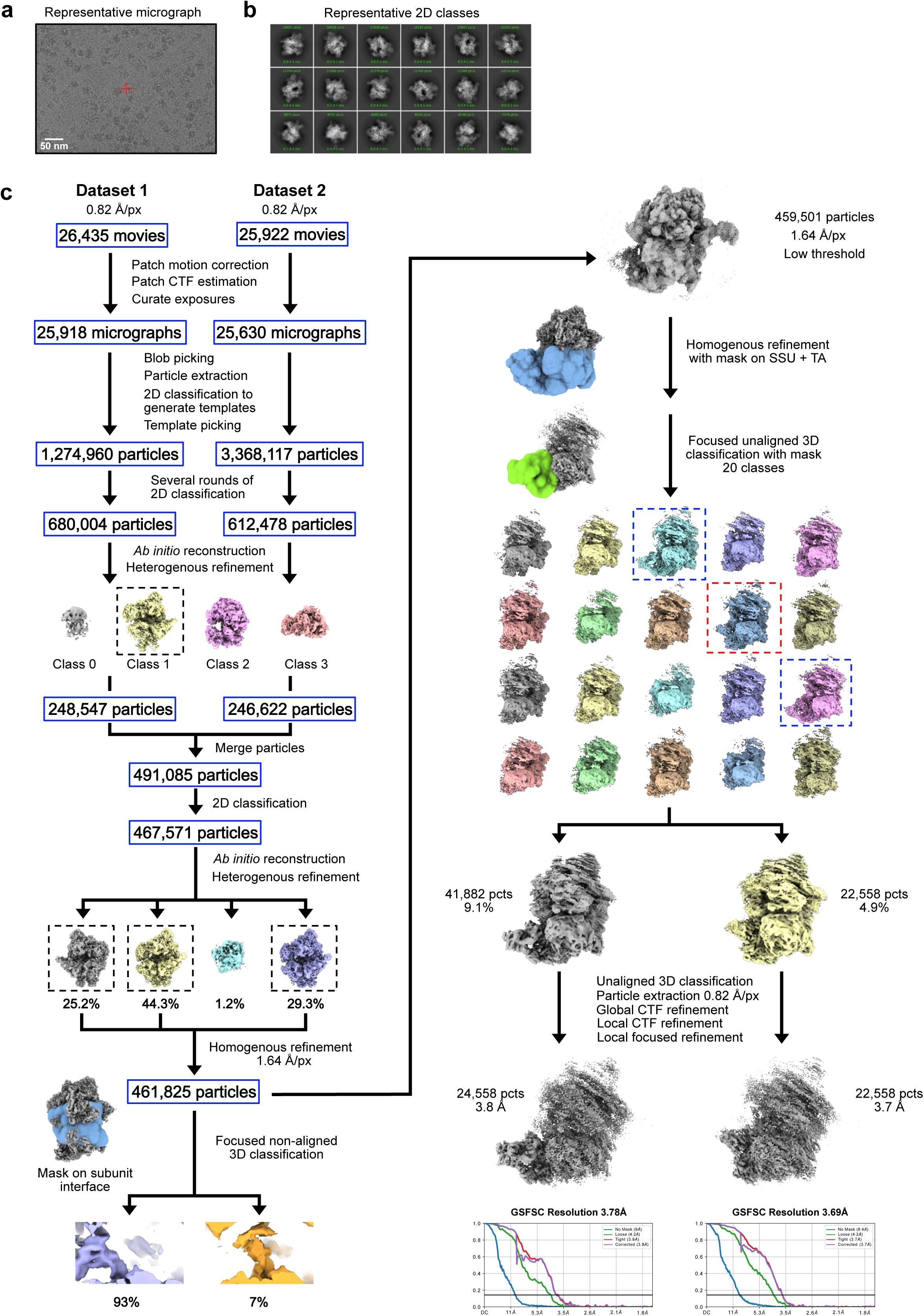
Cryo-EM processing workflow for *tuf1Δ* mitoribosomes. **a,** Representative micrograph of mitoribosome particles from the *tuf1Δ* sample. **b,** Representative 2D class averages of mitoribosome particles from the *tuf1Δ* sample. **c,** Schematic depicting the overall cryo-EM data processing workflow for the stalled post-initiating mitoribosomes from the *tuf1Δ* sample. Two data sets collected from the same grid were subjected to similar pre-processing in cryoSPARC v4.3.1 containing Patch motion correction, Patch CTF estimation, particle picking, 2D classification, *ab-initio* reconstruction and heterogenous refinement before merging of particles. Merged mitoribosome monosome particles were subsequently subjected to homogenous refinement followed by focused non-aligned 3D classification with a binary mask covering the subunit interface (to investigate tRNA occupancy) or the mitoribosomal SSU and the mRNA exit channel. The latter resulted in two major classes with additional densities located at the mRNA exit channel. Local refinement of the two classes resulted in estimated resolutions of 3.8 Å and 3.7 Å, respectively.

**Extended Data Figure 7:**
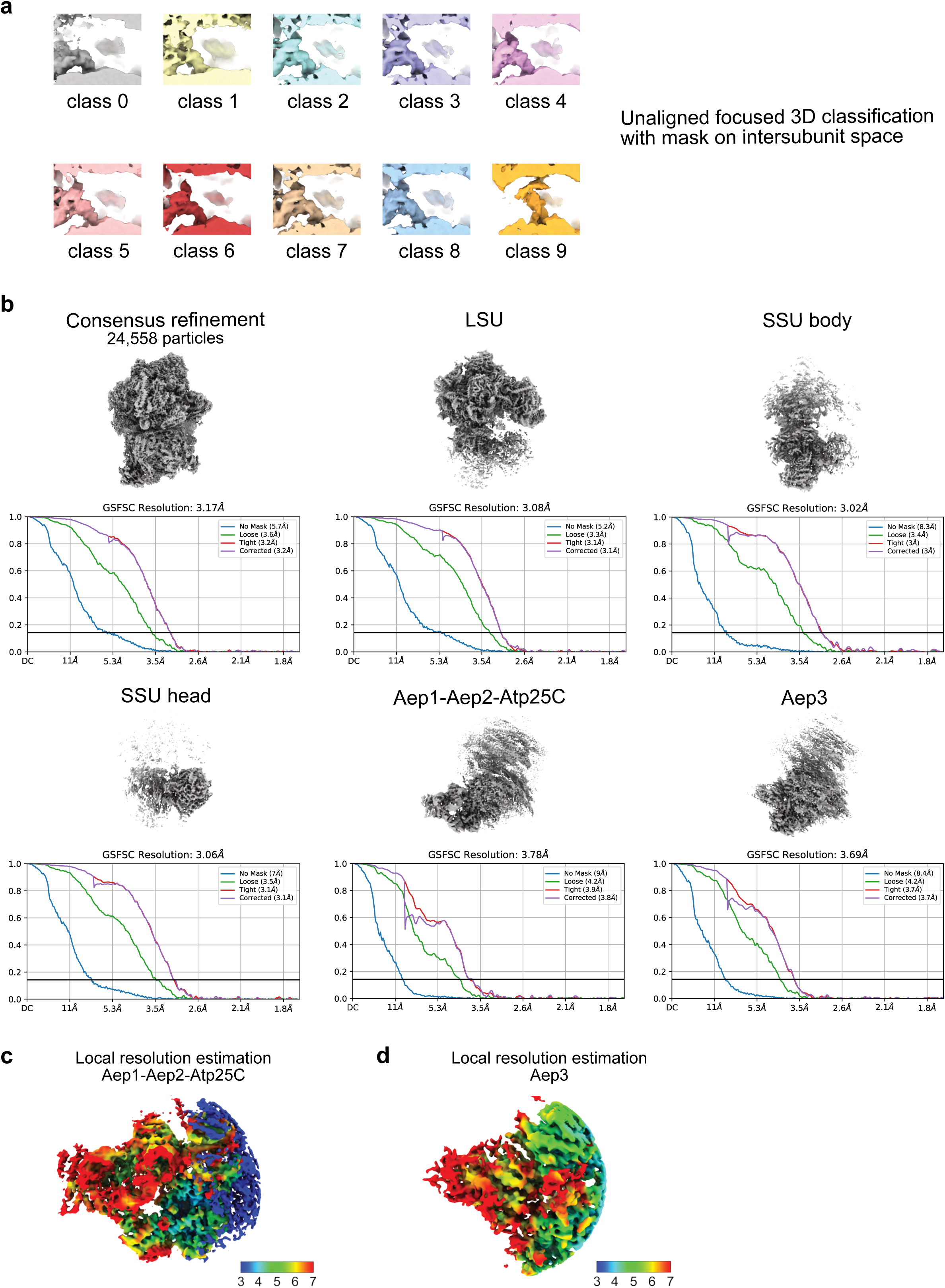
tRNA occupancy, refined maps and local resolution estimations for the *tuf1Δ* mitoribosomes. **a,** Visualization of the 10 classes after focused 3D classification using a binary mask on the subunit interface. Classes 0 to 8 show a tRNA density in the E-site, interacting with the L7/12 stalk of the LSU, while solely class 9 showed a density occupying the P-site. **b,** Density map of the consensus refinement using 24,558 particles and locally refined density maps of the LSU, SSU body, SSU head, Aep1-Aep2-Atp25C and Aep3 and their respective gold-standard Fourier shell correlation (GSFSC) curves with a cut-off at 0.143. **c,** Local resolutions of the Aep1-Aep2-Atp25C and Aep3 focused refined maps, with local resolution estimations ranging between 4 and 7 Å for the protein subunits and 6 Å and above for the mRNA densities.

**Extended Data Figure 8:**
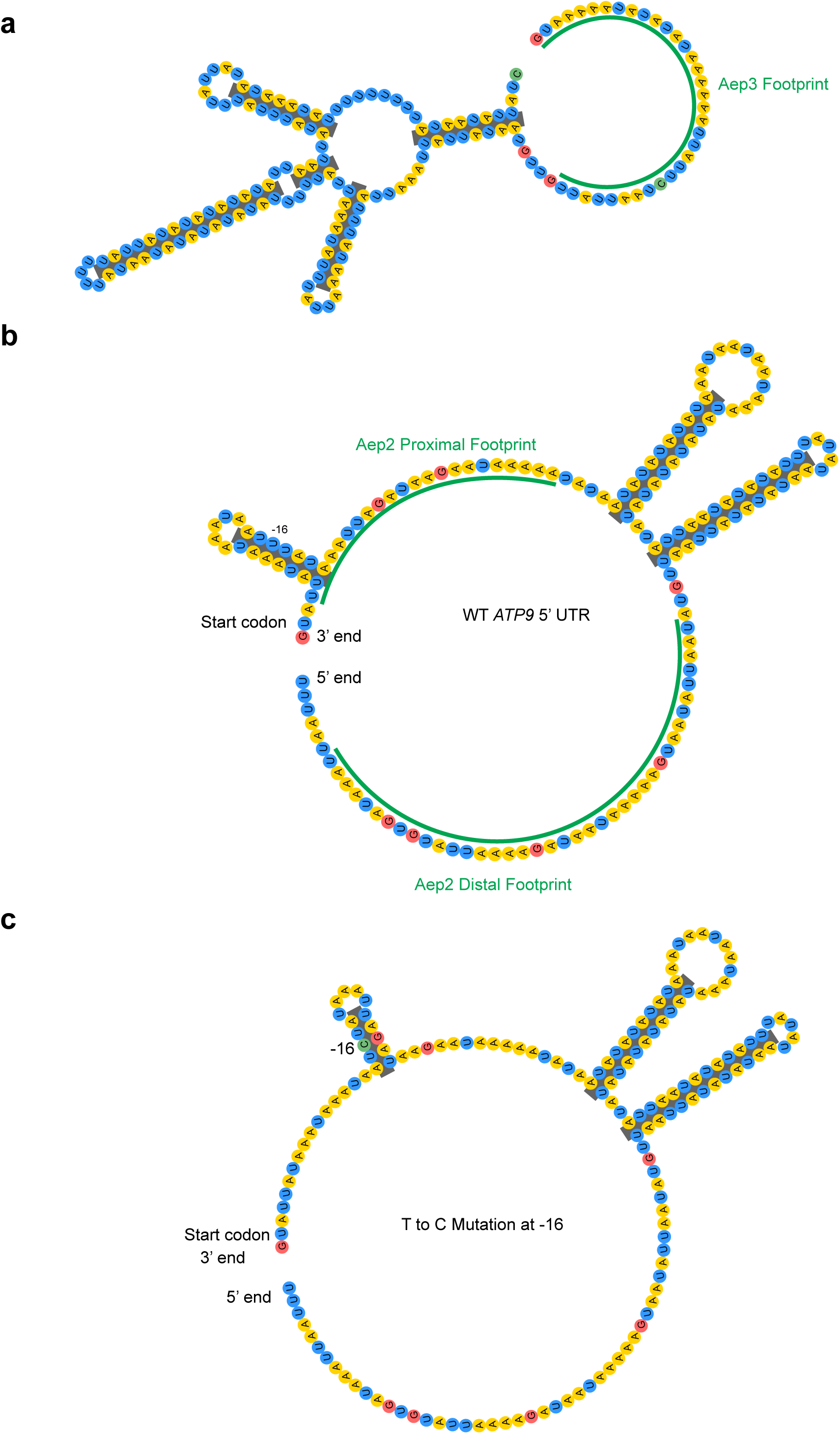
Predicted secondary structures in the 5’ UTRs of *ATP8* and *ATP9*. **a,** Eternafold secondary structure prediction of the 5’ UTR of *ATP8* upstream of the Aep3-mitoribosome footprint (underlined in green). **b,** Eternafold was used to predict the secondary structure of the 5’ UTR of *ATP9* from the start codon until just upstream of the distal Aep2 sel-mitoRP footprint. Both the proximal and distal Aep2 footprints are underlined in the predicted RNA structure diagram. A stem loop is predicted to form just upstream of the start codon where we see an overlapping Aep2 footprint. Two larger stem loops are predicted to form between the two binding sites, bringing the two footprints in close proximity in three-dimensional space. **c,** A mutation known to rescue a temperature sensitive mutant of Aep2 in which a T at -16 is converted to a C was modeled in Eternafold to visualize how the secondary structure of the RNA might change. A novel stem loop is generated due to base pairing between the C at -16 and the G at -26.

**Extended Data Figure 9:**
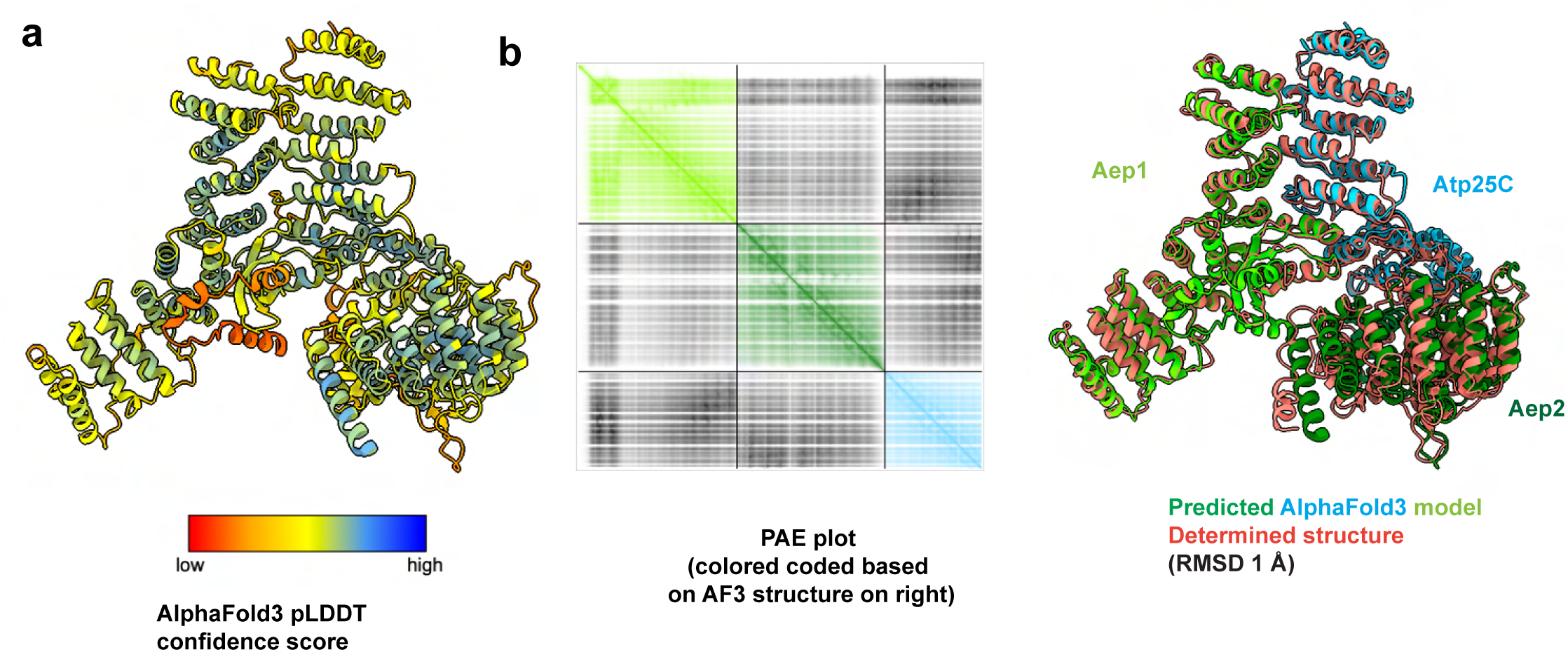
AlphaFold modeling of *ATP9* TA Complex. **a,** AlphaFold3 model of Aep1-Aep2-Atp25C color coded based on the pLDDT confidence score. **b,** PAE plot showing the Aep1-Aep2-Atp25C complex predicted by AlphaFold3 (left) color-coded to match the structure shown on the right. Overlay of the color-coded Aep1-Aep2-Atp25C complex obtained by AlphaFold3 with the structure obtained by cryo-EM shown in salmon red (right).

**Extended Data Figure 10:**
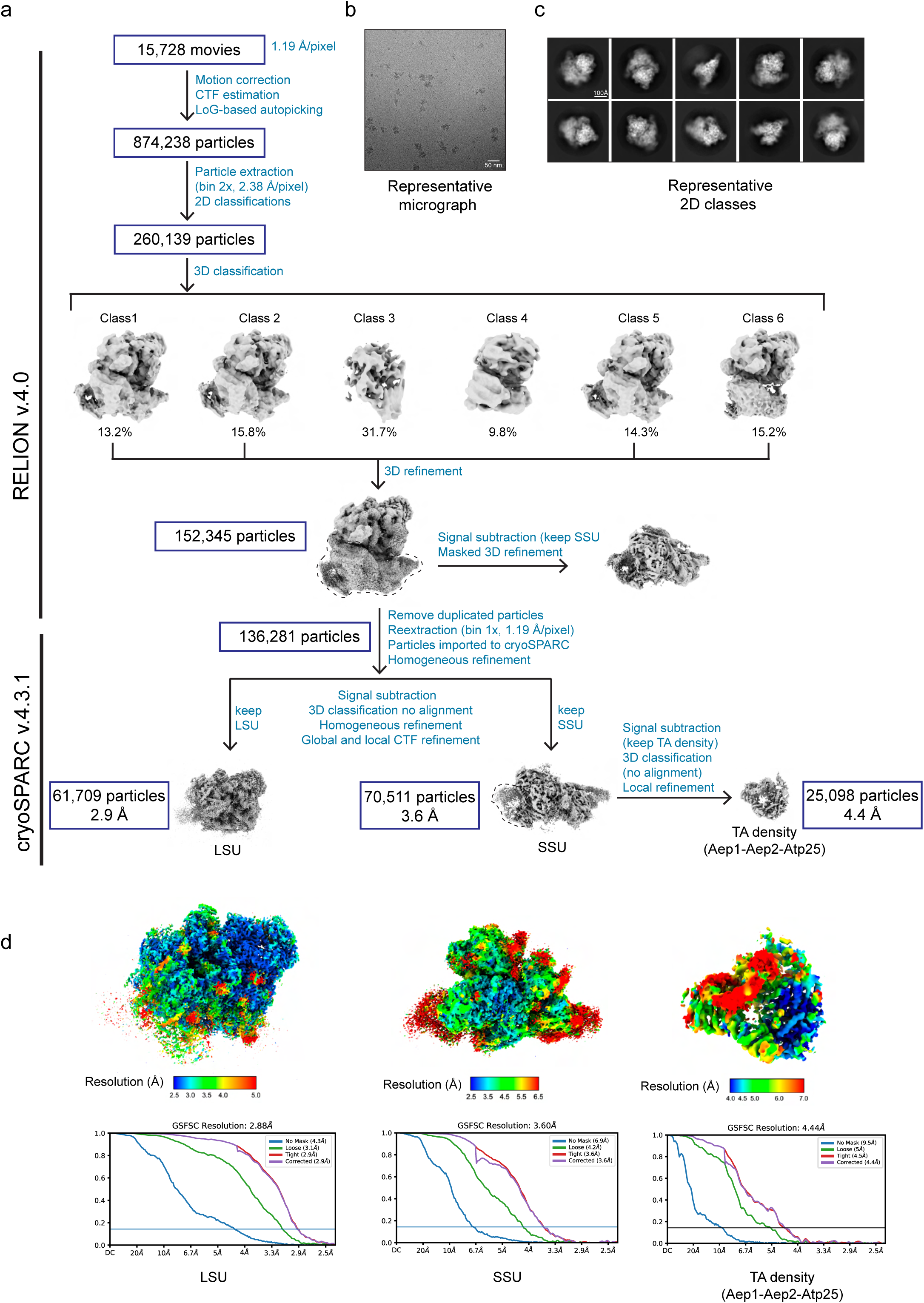
Cryo-EM data processing for direct purification of Aep2-bound mitoribosomes. **a,** Flow chart showing details of cryo-EM data processing of Aep2-bound mitoribosomes in RELION v4.0 and cryoSPARC v.4.3.1, respectively. **b,** A representative micrograph is shown with scale bar corresponding to 50 nm. **c,** Representative 2D classes of the Aep2-affinity purified mitoribosomes. **d,** Local resolution estimation (color coded to depict the local resolution of different parts) and gold-standard Fourier shell correlation (GSFSC) plot for each focused-refined map (FSC cut-off 0.143).

## Notes

### Competing Interest Statement

The authors have declared no competing interest.

