## Supplementary Tables for "Translational activators align mRNAs at the small mitoribosomal subunit for translation initiation"

**Supplementary Table 1.** Cryo-EM data collection, processing, refinement and validation statistics

|  | **Tuf1 KO Dataset 1** | **Tuf1 KO Dataset 2** | | |
| --- | --- | --- | --- | --- |
| **Data collection and processing** | | | | |
| Microscope | FEI Titan Krios | | | FEI Titan Krios |
| Camera | Gatan K3 | | | Gatan K3 |
| Magnification | 165,000 | | | 165,000 |
| Voltage (kV) | 300 | | | 300 |
| Electron exposure (e^-^/Å^2^) | 38 | | | 38 |
| Defocus range (μm) | -0.4 to -2.6 | | | -0.4 to -2.6 |
| Pixel size (Å) | 0.828 | | | 0.828 |
| Symmetry imposed | C1 | | | C1 |
| Number of images | 26,918 | | | 25,922 |
| Initial particles (no.) | 1,274,960 | | | 3,368,117 |
| Merged particles (no.) | 459,501 | | | |
| **Refinement** | | | | |
|  | **Aep1-Aep2-Atp25C** | | | **Aep3** |
| Refined particles (no.) | 41,190 | | | 22,558 |
| FSC threshold | 0.143 | | | 0.143 |
| Map resolution (Å) (consensus/LSU/SSU body/ SSU head/TA complex) | 3.03/2.93/2.96/ 2.93/3.76 | | | 2.96/2.82/3.02/ 3.01/3.69 |
| Map sharpening *B* factor (Å^2^) (consensus/LSU/SSU body/ SSU head/TA complex) | -41/-47/-36/ -40/-50 | | | -25/-30/-25/ -30/-30 |
| **Model composition** | | | | |
|  | **Aep1-Aep2-Atp25C** | | **Aep3** | |
| Initial model (PDB code) | 5MRC, 8OM4,  Alphafold2 | | 5MRC, 8OM4,  Alphafold2 | |
| Non-hydrogen protein atoms | 220,346 | | 214,164 | |
| Protein residues | 15,702 | | 15,088 | |
| Nucleotides | 4401 | | 4345 | |
| Protein chains | 76 | | 74 | |
| RNA chains | 4 | | 4 | |
| **Validation** |  | |  | |
| RMSD (bonds) | | | | |
| Length (Å) | 0.004 | | 0.004 | |
| Angles (°) | 0.528 | | 0.606 | |
| Molprobity score | 1.71 | | 1.71 | |
| Clashscore | 7.61 | | 7.45 | |
| Rotamer outliers (%) | 0.03 | | 0.04 | |
| Ramachandran plot | | | | |
| Favored (%) | 95.75 | | 95.64 | |
| Allowed (%) | 4.25 | | 4.36 | |
| Outliers (%) | 0.00 | | 0.00 | |

**Supplementary Table 2**. Cryo-EM data collection and processing of Aep2-bound mitoribosome

| **Data collection and processing** | | | |
| --- | --- | --- | --- |
| Microscope | FEI Titan Krios | | |
| Camera | Falcon 4i | | |
| Magnification | 105,000 | | |
| Voltage (kV) | 300 | | |
| Electron exposure (e–/Å^2^) | 48.5 | | |
| Defocus range (μm) | -0.8 to -2.0 | | |
| Pixel size (Å) | 0.825 | | |
| Symmetry imposed | C1 | | |
| Number of images | 15,728 | | |
| Initial particles picked (no.) | 874,238 | | |
|  | **LSU** | **SSU** | **Aep1-Aep2-Atp25C** |
| Final particles (no.) | 61,709 | 70,511 | 25,098 |
| Map resolution (Å)  FSC threshold | 2.9  0.143 | 3.6  0.143 | 4.4 0.143 |
| Map resolution range (Å) | 2.5-5.0 | 2.5-6.5 | 4.0-7.0 |

**Supplementary Table 3.** Yeast strains used in this study

| **Strain** | **Description** | **Background** | **Genotype** | **Source** |
| --- | --- | --- | --- | --- |
| YSC052 | Mrps17-FLAG | S288C | *mrps17::MRPS17-3XF* *MAT*a *GAL2 HAP1* *MIP1[S]* G1981A *ura3Δ0* | Couvillion et al., 2016^1^ |
| YSC058 | Mrps17-HA | S288C | *mrps17::MRPS17-3XHA* *MAT*a *GAL2 HAP1* *MIP1[S]* G1981A *ura3Δ0* | Couvillion et al., 2016^1^ |
| YSC353 | Aep1-FLAG; Mrps17-HA | S288C | *mrps17::MRPS17-3XHA aep1::AEP1-3XF* *MAT*a *GAL2 HAP1* *MIP1[S]* G1981A *ura3Δ0* | This study |
| YSC355 | Aep2-FLAG; Mrps17-HA | S288C | *mrps17::MRPS17-3XHA aep2::AEP2-3XF* *MAT*a *GAL2 HAP1* *MIP1[S]* G1981A *ura3Δ0* | This study |
| YSC359 | Aep2-Myc; Mrps17-FLAG; Img1-HA | S288C | *mrps17::MRPS17-3XF aep2::AEP2-3XMyc img1::IMG1-3XHA* *MAT*a *GAL2 HAP1* *MIP1[S]* G1981A *ura3Δ0* | This study |
| YSC362 | Aep1-Myc; Aep2-FLAG; Mrps17-HA | S288C | *mrps17::MRPS17-3XHA aep1::AEP1-3XMyc aep2::AEP2-3XF* *MAT*a *GAL2 HAP1* *MIP1[S]* G1981A *ura3Δ0* | This study |
| YSC363 | Aep1-FLAG; Aep2-Myc; Mrps17-HA | S288C | *mrps17::MRPS17-3XHA aep1::AEP1-3XF aep2::AEP2-3XMyc* *MAT*a *GAL2 HAP1* *MIP1[S]* G1981A *ura3Δ0* | This study |
| YSC364 | Atp22-FLAG; Mrps17-HA | S288C | *mrps17::MRPS17-3XHA atp22::ATP22-3XF MAT*a *GAL2 HAP1* *MIP1[S]* G1981A *ura3Δ0* | This study |
| YSC367 | Cbp1-FLAG; Mrps17-HA | S288C | *mrps17::MRPS17-3XHA cbp1::CBP1-3XF MAT*a *GAL2 HAP1* *MIP1[S]* G1981A *ura3Δ0* | This study |
| YSC369 | Pet309-FLAG; Mrps17-HA | S288C | *mrps17::MRPS17-3XHA pet309::PET309-3XF MAT*a *GAL2 HAP1* *MIP1[S]* G1981A *ura3Δ0* | This study |
| YSC370 | Aep3-FLAG; Mrps17-HA | S288C | *mrps17::MRPS17-3XHA aep3::AEP3-3XF MAT*a *GAL2 HAP1* *MIP1[S]* G1981A *ura3Δ0* | This study |
| YSC371 | Pet111-FLAG; Mrps17-HA | S288C | *mrps17::MRPS17-3XHA pet111::PET111-3XF MAT*a *GAL2 HAP1* *MIP1[S]* G1981A *ura3Δ0* | This study |

**Supplementary Table 4.** Plasmids used in this study

| **Plasmid** | **Use** | **Source** |
| --- | --- | --- |
| p3xFs-URA3-3xFs | Scarless C-terminal FLAG-tagging | Couvillion et al., 2016^1^ Modified from Moqtaderi and Struhl, 2008^2^ |
| pMPY-3xHA | Scarless C-terminal HA-tagging | Schneider, B. L., et al.,1995^3^ |
| pMPY-3xMyc | Scarless C-terminal Myc-tagging | Schneider, B. L., et al.,1995^3^ |
| YEplac181-RNR1 | Expression of Rnr1 in *TUF1* deletion strain | Zeng et al., 2018^4^ |
| pRS316-VAR1 | Expression of Var1 in *TUF1* deletion strain | Zeng et al., 2018^4^ |
| pFA6-KanMX4 | Deletion of *TUF1* | Wach et al., 1994^5^ |

**Supplementary Table 5.** Primers used in this study

| **Name** | **Sequence** | **Use** | **Source** |
| --- | --- | --- | --- |
| JB010 | GAAATATTTGAATTGCACTGAACGAGAAGCTTTACGCCCAAGGGAACAAAAGCTGG | Forward primer 3’ Myc/FLAG tag for Aep1 | This study |
| JB011 | TGAGTTGCTCTTATGCTGCCTTTTTTTTAATATAGACGATCTATAGGGCGAATTGG | Reverse primer 3’ Myc/FLAG tag for Aep1 | This study |
| JB014 | TGACGATGAAGATGATGGTATGATTATAGGTAGCCTTTGGAGGGAACAAAAGCTGG | Forward primer 3’ Myc/FLAG tag for Aep2 | This study |
| JB015 | ATATGGAGAAAAATAAATGCTAATTCTACTTACACTTTTCCTATAGGGCGAATTGG | Reverse primer 3’ Myc/FLAG tag for Aep2 | This study |
| JB034 | ATGTACAAATATTATAAGAGAGACGTTGAAAAGTCTAAATAGGGAACAAAAGCTGG | Forward primer 3’ FLAG tag for Atp22 | This study |
| JB035 | ATGAATGAATATTTACTATTTACTAGTGCTCATCTGGATACTATAGGGCGAATTGG | Reverse primer 3’ FLAG tag for Atp22 | This study |
| JB038 | CAAAAAAGAGCTAGTAAAGAGGAGAATAGTTGGGGAGGTTAGGGAACAAAAGCTGG | Forward primer 3’ FLAG tag for Aep3 | This study |
| JB039 | AAATTCCTGTCCAAATGCCAGGGCACTTATGCCTCCCTCACTATAGGGCGAATTGG | Reverse primer 3’ FLAG tag for Aep3 | This study |
| JB073 | TAAAAAGCATGGTGTGTCGGCTGTCAAACGTTACTTAAGAAGGGAACAAAAGCTGG | Forward primer 3’ FLAG tag for Cbp1 | This study |
| JB074 | TTTGCTTTGTTATTTATATCGTAAATGTGCGTTTGGCCGTCTATAGGGCGAATTGG | Reverse primer 3’ FLAG tag for Cbp1 | This study |
| JB077 | ACTGAGGAAATCTAAGAGAGTATTACCTGTGAGTAAATTCAGGGAACAAAAGCTGG | Forward primer 3’ FLAG tag for Pet309 | This study |
| JB078 | ATATATATGCAGTTGATTATACAATATGATATATGCATTTCTATAGGGCGAATTGG | Reverse primer 3’ FLAG tag for Pet309 | This study |
| JB081 | GAAAATGAAGCTTTTTGAAGAGAATAAAAAGGAGGAGGAGAGGGAACAAAAGCTGG | Forward primer 3’ FLAG tag for Pet111 | This study |
| JB082 | ATTTACACGTGAGAGAAAGGAAGGTAAATAACTGAAAAGACTATAGGGCGAATTGG | Reverse primer 3’ FLAG tag for Pet111 | This study |
| TUF1 KO forward primer | CTATTTTGTGCTTTCAGTTTTATTCTAGCTCGACAAAGGTAACAGACAAAACCAGCTGAAGCTTCGTACGCTGC | Forward primer for generating *TUF1* deletion cassette | This study |
| TUF1 KO reverse primer | GGGGAAATGAACAGAATATATAGAAATATACTCCAGTTGCATCAATAAGTCCGCATAGGCCACTAGTGGATCTG | Reverse primer for generating *TUF1* deletion cassette | This study |
